# African Swine Fever Virus and host response - transcriptome profiling of the Georgia 2007/1 strain and porcine macrophages

**DOI:** 10.1101/2021.07.26.453801

**Authors:** Gwenny Cackett, Raquel Portugal, Dorota Matelska, Linda Dixon, Finn Werner

## Abstract

African swine fever virus (ASFV) has a major global economic impact. With a case fatality in domestic pigs approaching 100%, it currently presents the largest threat to animal farming. Although genomic differences between attenuated and highly virulent ASFV strains have been identified, the molecular determinants for virulence at the level of gene expression have remained opaque. Here we characterise the transcriptome of ASFV genotype II Georgia 2007/1 (GRG) during infection of the physiologically relevant host cells, porcine macrophages. In this study we applied Cap Analysis Gene Expression sequencing (CAGE-seq) to map the 5’ ends of viral mRNAs at 5 and 16 hours post-infection. A bioinformatics analysis of the sequence context surrounding the transcription start sites (TSSs) enabled us to characterise the global early and late promoter landscape of GRG. We compared transcriptome maps of the GRG isolate and the lab-attenuated BA71V strain that highlighted GRG virulent-specific transcripts belonging to multigene families, including two predicted MGF 100 genes I7L and I8L. In parallel, we monitored transcriptome changes in the infected host macrophage cells. Of the 9,384 macrophage genes studied, transcripts for 652 host genes were differentially regulated between 5 and 16 hours-post-infection compared with only 25 between uninfected cells and 5 hours post-infection. NF-kB activated genes and lysosome components like S100 were upregulated, and chemokines such as CCL24, CXCL2, CXCL5 and CXCL8 downregulated.

**Importance:** African swine fever virus (ASFV) causes haemorrhagic fever in domestic pigs with case fatality rates approaching 100%, and no approved vaccines or antivirals. The highly-virulent ASFV Georgia 2007/1 strain (GRG) was the first isolated when ASFV spread from Africa to the Caucasus region in 2007. Then spreading through Eastern Europe, and more recently across Asia. We used an RNA-based next generation sequencing technique called CAGE-seq to map the starts of viral genes across the GRG DNA genome. This has allowed us to investigate which viral genes are expressed during early or late stages of infection and how this is controlled, comparing their expression to the non-virulent ASFV-BA71V strain to identify key genes that play a role in virulence. In parallel we investigated how host cells respond to infection, which revealed how the ASFV suppresses components of the host immune response to ultimately win the arms race against its porcine host.

## Introduction

ASFV originated in Sub-Saharan Africa where it remains endemic. However, following the introduction in 2007, of a genotype II isolate to Georgia (1), and subsequent spread in Russia and Europe. The virus was then introduced to China in 2018 (2), from here it spread rapidly across Asia, strongly emphasizing this disease as a severe threat to global food security. ASFV is the only characterised member of the Asfarviridae family (3) in the recently classified Nucleocytoviricota (ICTV Master Species List 2019.v1) phylum (4, 5). ASFV has a linear double-stranded DNA (dsDNA) genome of ∼170–193 kbp encoding ∼150–∼200 open reading frames (ORFs). Until recently, little was known about either the transcripts expressed from the ASFV genome or the mechanisms of ASFV transcription. Much of what is known about transcription is extrapolated from vaccinia virus (VACV), a distantly-related Nucleocytoviricota member, from the Poxviridae family (6). ASFV encodes a eukaryotic-like 8-subunit RNA polymerase (RNAP), an mRNA capping enzyme and poly-A polymerase, all of which are carried within mature virus particles (7). These virions are transcription competent upon solubilisation in vitro (8) and support mRNA modification by including a 5’-methylated cap and a 3’ poly-adenylated (polyA) tail of ∼33 nucleotide-length (8, 9).

Viral genes are typically classified according to their temporal expression patterns - ASFV genes have historically been categorised as ‘immediate early’ when expressed immediately following infection, as ‘early genes’ following the onset of viral protein synthesis, as ‘intermediate genes’ after the onset of viral DNA replication, or as ‘late genes’ thereafter. The temporal regulation of transcription is likely enabled by different sets of general transcription initiation factors that recognise distinct early or late promoter motifs (EPM or LPM, respectively), as we previously investigated in the ASFV-BA71V strain (10), and address further in this study. EPM recognition is likely enabled by the ASFV homologue of heterodimeric VACV early transcription factor (VETF), consisting of D1133L (D6) and G1340L (A7) gene products, which bind the Poxvirus early gene promoter motif (11–13), which the ASFV EPM strongly resembles. Both ASFV-D6 and ASFV-A7 are late genes, i.e. synthesised late during infection (10) and packaged into virus particles (7). The ASFV LPM is less well defined than the EPM, but a possible initiation factor involved in its recognition is the ASFV-encoded viral homolog of the eukaryotic TATA-binding protein (TBP), expressed during early infection (10). By analogy with the VACV system, additional factors including homologs of A1, A2 and G8 may also contribute to late transcription initiation (6) We have recently carried out a detailed and comprehensive ASFV whole genome expression analysis using complimentary next-generation sequencing (NGS) results and computational approaches to characterise the ASFV transcriptome following BA71V infection of Vero cells at 5 hpi and 16 hpi post-infection (hpi) (10). Most of our knowledge about the molecular biology of ASFV, including gene expression, has been derived from cell culture-adapted, attenuated virus strains, such as BA71V infecting Vero tissue culture cells (9, 10). These model systems provide convenient models to study the replication cycle but have deletions of many genes that are not essential for replication, but have important roles in virulence within its natural porcine hosts. (14–16). To date 24 ASFV genotypes have been identified in Africa (16–23), while all strains spreading across Asia and Europe belong to the Type II genotype. Most of these are highly virulent in domestic pigs and wild boar, including the ASFV Georgia 2007/1 (GRG) (24), and the Chinese ASFV Heilongjiang, 2018 (Pig/HLJ/18) (25) isolates. Though a number of less virulent isolates have been identified in wild boar in the Baltic States and domestic pigs in China (26–29). It is crucial to understand the similarities and commonalities between ASFV strains, and to characterise the host response to these in order to understand the molecular determinants for ASFV pathogenicity. Information about the gene content and genome organisation can be gained from comparing virus genome sequences. However, only functional genomics such as transcriptome or proteome analyses can provide information about the differences in gene expression programmes and the host responses to infection.

On the genome level, most differences between virulent (e.g. GRG) and attenuated (e.g. lab-attenuated BA71V) ASFV strains reside towards the genome termini. Figure 1a shows a whole genome comparison of GRG (left) and BA71V (right) strains with the sequence conservation colour coded in different shades of blue. The regions towards the ends of the genome are more dynamic compared to the central region which is highly conserved, as genes at the termini are prone to deletion, duplication, insertion, and fusion (17, 30). Most of the GRG-specific genes are expressed early during infection (early genes are colour coded blue in the outer arch of Figure 1a) and many belong to Multi-Gene Families (MGFs, purple in the inner arch). The functions of many MGF members remain poorly understood, though variation among MGFs is linked to virulence (31) and deleting members of MGF 360 and 505 families has been shown to reduce virulence (32, 33). Deletion of MGF 505-7R or MGF 110-9L also partially attenuated the virus in pigs (34, 35). In contrast deletion of MGF 110-1L and MGF 100-1R did not reduce virus virulence (17). Members of MGF 110 are highly expressed both on the mRNA and protein level in infections with the BA71V isolate or OURT88/3 (10, 36), suggesting MGF 110 holds importance during infection. Overall, the functions of MGF 360 and 505 members are better characterised than other MGFs, playing a role in evading the host type I interferon (IFN) response (15,32,37–40). In summary, comparing the expression of ASFV genes, especially MGFs, between the virulent GRG- and the lab adapted BA71V strains, is fundamental in identification of virulence factors and better MGF characterisation.

**Figure 1.**
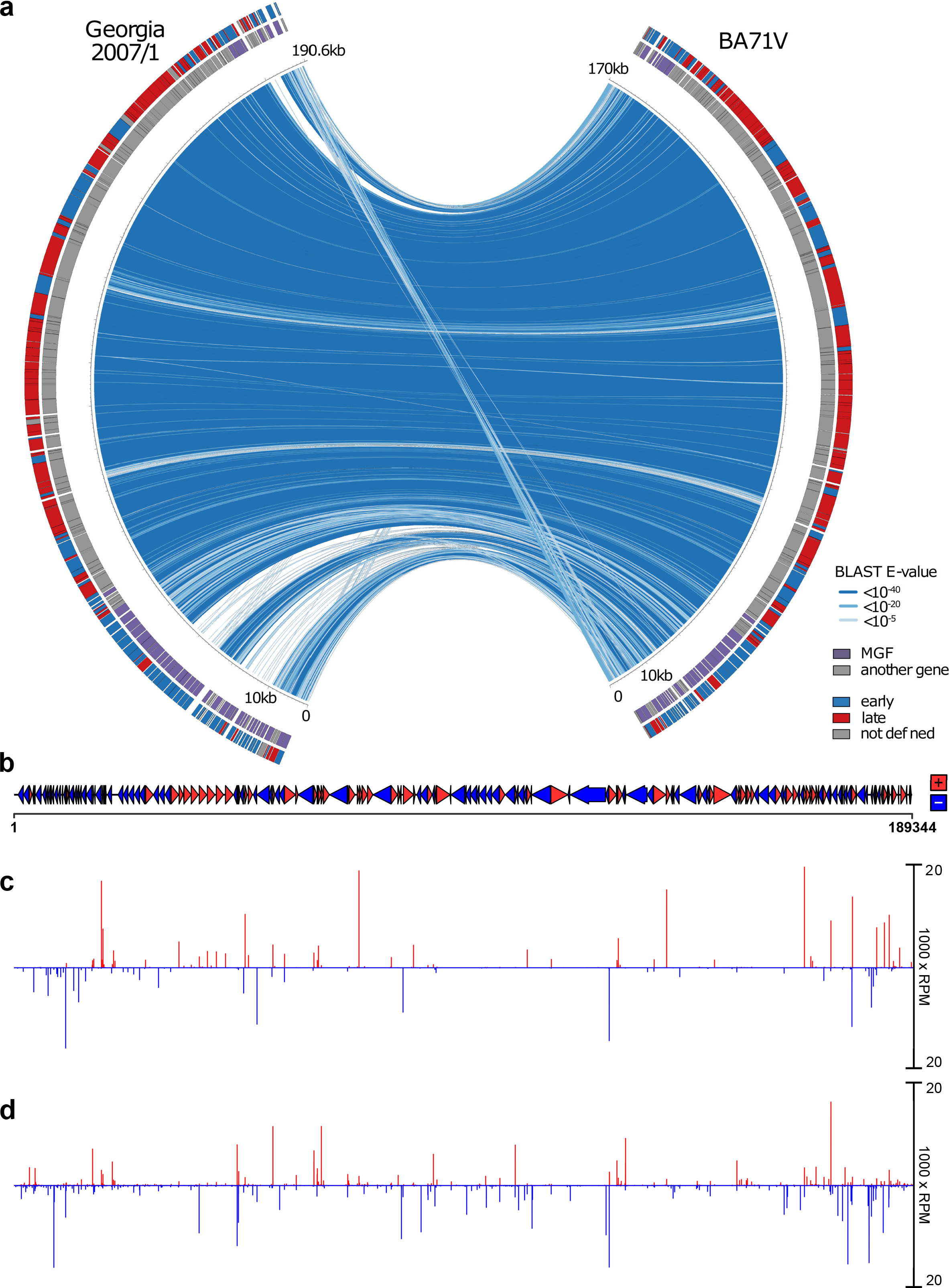
Functional genome annotation of ASFV GRG. (a) Comparison between the genomes of BA71V and GRG, generated with Circos (http://circos.ca/). Blue lines represent sequence conservation (Blast E-values per 100 nt). The Inner ring represents genes defined as MGF members (purple), and all others (grey). The outer ring shows annotated genes which we have defined as early or late according to downregulation or upregulation between 5 hpi and 16 hpi from DESeq2 analysis. (b) 189 GRG annotated ORFs are represented as arrows and coloured according to strand. CAGE-seq peaks across the GRG genome at 5 hpi (c) and 16 hpi (d), normalized coverage reads per million mapped reads (RPM) of 5’ ends of CAGE-seq reads. The coverage was capped at 20000 RPM for visualisation, though multiple peaks exceeded this. DeepTools (112), was used to convert bam files to bigwig format and imported into Rstudio for visual representation via packages ggplot, ggbio, rtracklayer, and gggenes was used to generate the ORF map in (b).

Macrophages are the primary target cells for ASFV, they are important immune effector cells that display remarkable plasticity allowing efficient response to environmental signals (41). There are some studies which have investigated how host macrophages respond to infection, including a microarray analysis of primary swine macrophage cells infected with virulent GRG (42). There are two RNA-seq studies of whole blood or tissues isolated from pigs post-mortem, which were infected with either a low pathogenic ASFV-OURT 88/3 or ASFV-GRG (43), or infected with a pathogenic Chinese isolate ASFV-SY18 (44). Recently, two reports have been published about the transcriptomic response of porcine macrophages to infection with a virulent Chinese genotype II isolate using a low multiplicity of infection (MOI: 1) and classical RNA-seq (45, 46), but due to different experimental conditions the varying results are somewhat challenging to compare with other studies. It must also be remembered that neither these classical RNA-seq nor microarray analyses, have sufficient resolution to accurately capture viral gene expression in the compact ASFV genome alongside that of the host.

Here we applied CAGE-seq to characterise the transcriptome of the highly virulent GRG isolate (24), in primary porcine macrophages, the biologically relevant target cells for ASFV infection. In this study we used a high multiplicity of infection (MOI: 5), so that transcripts expressed during a single cycle time course could be measured without the complication of variable proportions of uninfected cells being present. We investigated the differential gene expression patterns of viral mRNAs at early and late time points of 5- and 16 hpi, and mapped the viral promoter motifs. Importantly, we have compared the expression levels and temporal regulation of genes conserved in both the virulent GRG isolate, and attenuated tissue-culture adapted BA71V strain. With a few exceptions, both mRNA expression levels and temporal regulation of the conserved genes are surprisingly similar. This confirms that it is not deregulation of their conserved genes, but the virulent isolate-specific genes, which are the key determinants for ASFV virulence. Most of these genes are MGF members, likely involved in suppression of the host immune-response. Indeed, transcriptome analysis of the porcine macrophages upon GRG infection reflects a modulation of host immune response genes, although the bulk of the ∼ 9000 genes studied did not significantly change expression levels during infection.

## Results

### Genome-wide Transcription Start Site-Mapping

We infected primary porcine alveolar macrophages with ASFV GRG at a high multiplicity of infection (MOI 5.0), isolated total RNA at 5 hpi and 16 hpi and sequenced using CAGE-seq (Supplementary Table 1a). The resulting mRNA 5’ ends were mapped to the GRG genome (Figure 1b) resulting in the annotation of 229 and 786 TSSs at 5 and 16 hpi, respectively (Figure 1c and d, from Supplementary Table 1b and c, respectively). The majority of TSSs were identified within 500 bp upstream of the start codon of a given ORF, a probable location for a *bona fide* gene TSS. The strongest and closest TSSs upstream of ORFs were annotated as ‘primary’ TSS (pTSS, listed in Supplementary Table 1d) and in this manner we could account TSS for 177 out of 189 GRG ORFs annotated in the FR682468.1 genome. TSSs signals below the threshold for detection included MGF_110-11L, C62L, and E66L, the remainder being short ORFs designated as ‘ASFV_G_ACD’, predicted solely from the FR682468 genome sequence (24). The E66L ORF was originally predicted from only the BA71V genome sequence, but likewise was undetectable with CAGE-seq (10), making its expression unlikely. Our TSS mapping identified novel ORFs (nORFs) downstream of the TSS, which were included in the curated GRG genome map (Supplementary Table 1d includes pTSSs of annotated ORFs and nORFs in gene feature file or ‘GFF’ format). In addition to ORF-associated TSSs, some were located within ORFs (intra-ORF or ioTSS), or in between them (inter-ORF TSS), and all detected TSSs are listed in Supplementary Table 1b-c.

### Expression of GRG genes during Early and Late Infection

Having annotated TSSs across the GRG genome, we quantified the viral mRNAs originating from pTSSs from CAGE-seq data, normalising against the total number of reads mapping to the ASFV genome (i. e. RPM or reads per million mapped reads per sample). We compared gene expression between early and late infection, and simplistically defined genes as ‘early’ or ‘late’ if they are significantly down- or upregulated (respectively), using DESeq2 (47). In summary, 165 of the 177 detectable genes were differentially expressed (adjusted p-value or padj < 0.05, Supplementary Table 1e). Those showing no significant change were D345L, DP79L, I8L, MGF_100-1R, A859L, QP383R, B475L, E301R, DP63R, C147L, and I177L. 87 of those 165 differentially expressed genes were significantly downregulated, thus representing the ‘early genes’, while 78 of the 165 genes were upregulated or ‘late genes’. The majority of MGFs were early genes, apart from MGF 505-2R, MGF 360-2L and MGF 100-1L (Figure 2a). Figure 2b shows the expression patterns of GRG-exclusively expressed genes, which we defined as only having a detectable CAGE-seq TSS in GRG, and not in BA71V (regardless of presence in the BA71V genome). These unsurprisingly, consist of many MGFs (19), all of which were early genes (Figure 2b), barring MGF 100-1L. In addition genes l9R, l10L and l11L and several of the newly annotated short ORFs were specific to GRG.

**Figure 2.**
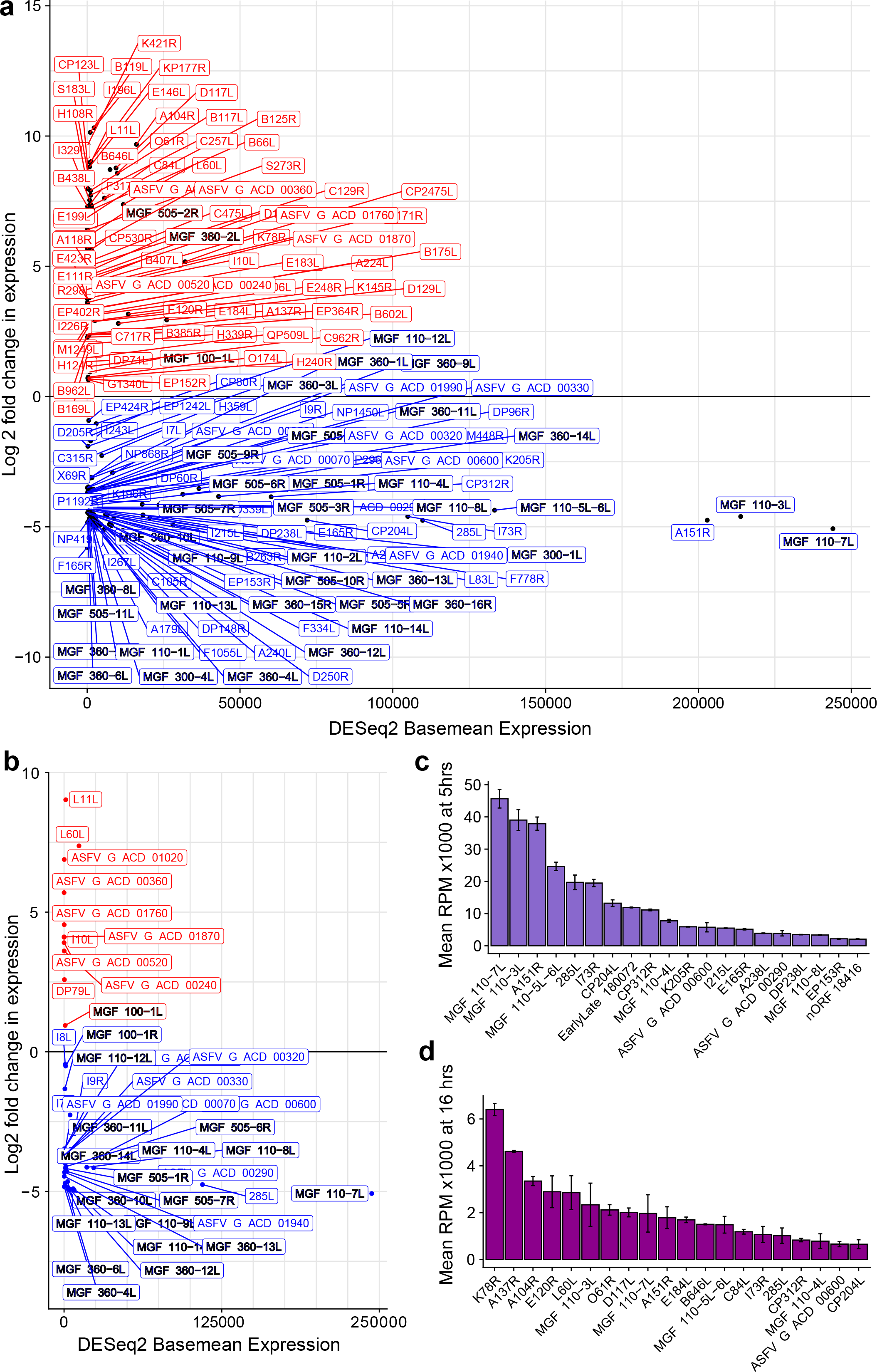
Summary of GRG gene expression (a) Expression profiles for 164 genes for which we annotated pTSSs from CAGE-seq and which showed significant differential expression. Log2 fold change and basemean expression values were from DESeq2 analysis of raw counts (see methods). Genes are coloured according to their log2 fold change in expression as red (positive: upregulated from 5 hpi to 16 hpi) or blue (negative: downregulated). MGFs are emphasised with a black outline to highlight their overrepresentation in the group of downregulated genes. (b) Expression profiles for 41 genes (excluding nORFs) only detected as being expressed in GRG and not BA71V, format as in (a). (c) Expression (RPM) of 20 highest-expressed genes at 5 hpi, error bars represent standard deviation between replicates. (d) Expression (RPM) of 20 highest-expressed genes at 16 hpi pi, error bars are the standard deviation between replicates.

We extracted the top twenty most highly expressed genes of GRG (as RPM) during 5 hpi (Figure 2c) and 16 hpi (Figure 2d) post-infection. Ten genes are shared between both top 20 lists: MGF 110-3L, A151R, MGF 110-7L, MGF 110-5L-6L, I73R, 285L, CP312R, ASFV_G_ACD_00600, MGF 110-4L, and CP204L. It is important to note that the relative expression values (RPM) for genes at 5 hpi are significantly higher than those at 16 hpi. This is consistent with our observations in the BA71V strain (10) and due to the increase in global viral transcript levels during late infection discussed below.

Supplementary Table 1f includes all the GRG annotated ORFs, their TSS locations during early and late infection, their relative distances if these TSS locations differ, and their respective 5’ Untranslated Region (UTR) lengths.

### GRG and BA71V Share Strong Similarity between Conserved Gene Expression

Next we carried out a direct comparison of mRNA levels from 132 conserved genes between the virulent GRG and attenuated BA71V (10) strain making use of our previously published CAGE-seq data. The relative transcript levels (RPM) of the genes conserved between the two strains showed a significant correlation at 5 hpi (Figure 3a) and 16 hpi (Figure 3b), supported by the heatmap in Supplementary Figure 1, the RPM for each gene, across both time-points and replicates, showing a strong congruence between the two strains. Of the 132 conserved genes, 125 showed significant differential expression in both strains. 119 of these 125 showed the same down- or up-regulated patterns of significant differential expression from 5 hpi to 16 hpi (Figure 3c, early genes in blue, late genes in red). The exceptions are D205R, CP80R, C315R, NP419L, F165R, and DP148R (MGF 360-18R), encoding RNA polymerase subunits RPB5 and RPB10 (15), Transcription Factor IIB (TFIIB) (15), DNA ligase (48), a putative signal peptide-containing protein, and a virulence factor (49), respectively. The ASFV-TFIIB homolog (C315R) is classified as an early gene in GRG but not in BA71V, in line with the predominantly early-expressed TBP (B263R), its predicted interaction partner. It is worth noting however, that D205R, CP80R, and C315R are close to the threshold of significance, with transcripts being detected at both 5 hpi and 16 hpi (Supplementary Table 1e).

**Figure 3.**
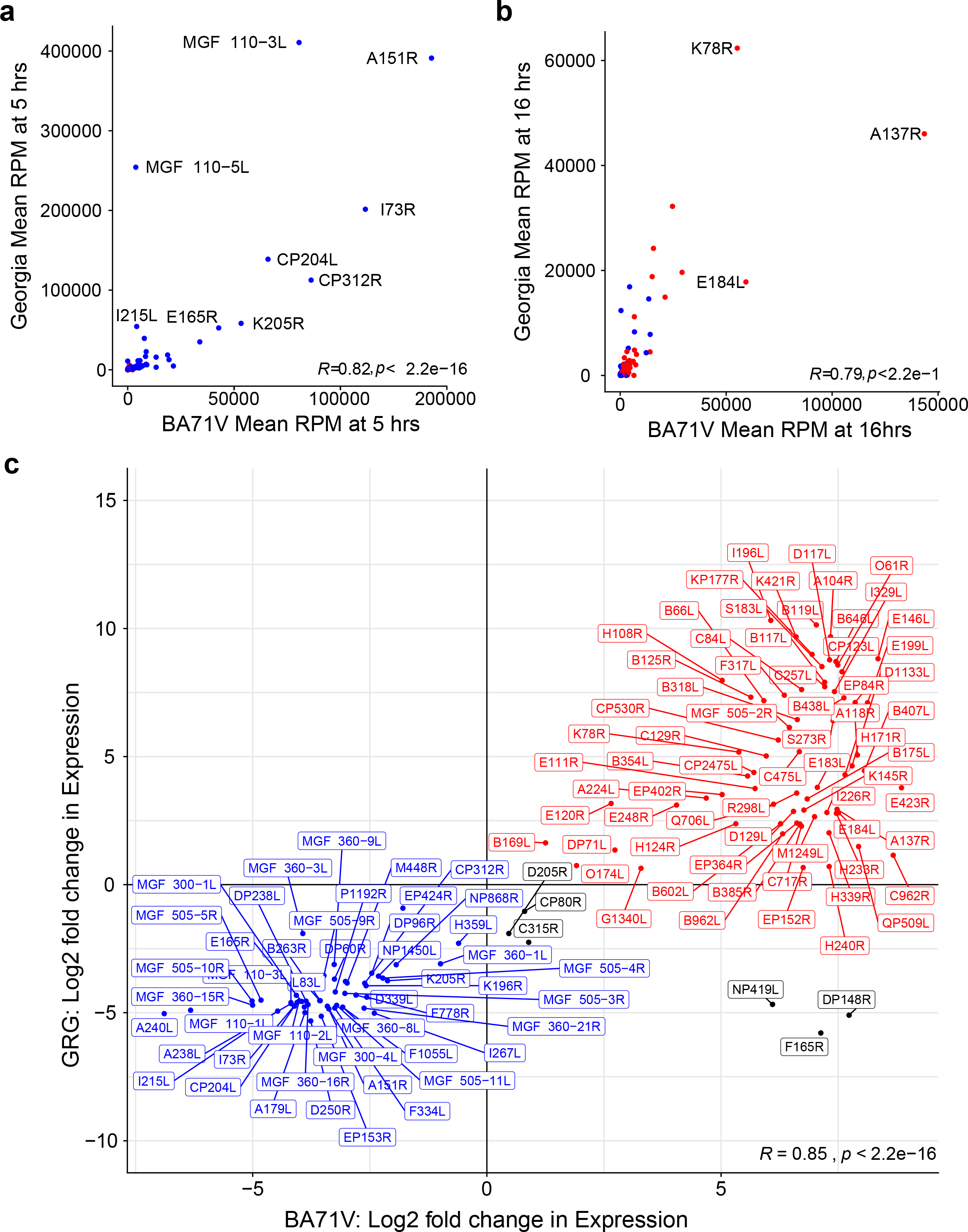
Comparison of gene expression profiles for genes shared between GRG and BA71V. Scatter plots of mean RPM across replicates for shared genes at 5 hpi (a) and 16 hpi (b), coloured according to whether genes show significant downregulation (blue), or upregulation (red) according to DESeq2 analysis in GRG. In both (b) and (c) genes with RPM values above 40000 RPM in either strain are labelled. (c) Comparison of log2 fold change in expression values of genes in GRG and BA71V, in blue are downregulated (early) genes in both strains, red are upregulated (late) genes in both strains, while the genes which disagree in their differential expression patterns between strains are in black. R represents the Pearson Correlation coefficient for each individual plot in (a), (b), and (c). Due to inconsistencies in their genome annotations, two genes were omitted from the BA71V-GRG transcriptome comparisons in Figures 2b and 3a-d: EP296R in GRG known as E296R in BA71V, and C122R (GRG) is the old nomenclature for C105R (BA71V), which are now correctly named in Supplementary Table 1e and Figure 2a. Both genes showed the same early expression patterns in BA71V (10) and GRG (Supplementary Table 1e) so would strengthen the patterns observed.

### Increased and pervasive transcription during late infection

During late infection of BA71V (10), we noted an increase in genome-wide mRNA abundance, as well as an increasing number of TSSs and transcription termination sites, reminiscent of pervasive transcription observed during late infection of Vaccinia virus (50). To quantify and compare the global mRNA increase both in BA71V and GRG, we calculated the ratio of read coverage per nucleotide, at 16 hpi versus 5 hpi (log2 transformed ratio of RPM), across the viral genome (Figure 4a, increase shown above- and decrease below the x-axis). This dramatic increase is due to the overall increase of virus mRNAs present, which is visible in both strains (Figure 4b), with a ∼2 fold increase in GRG from 5 hpi to 16 hpi, versus ∼8 fold in BA71V (Figure 4c).

**Figure 4.**
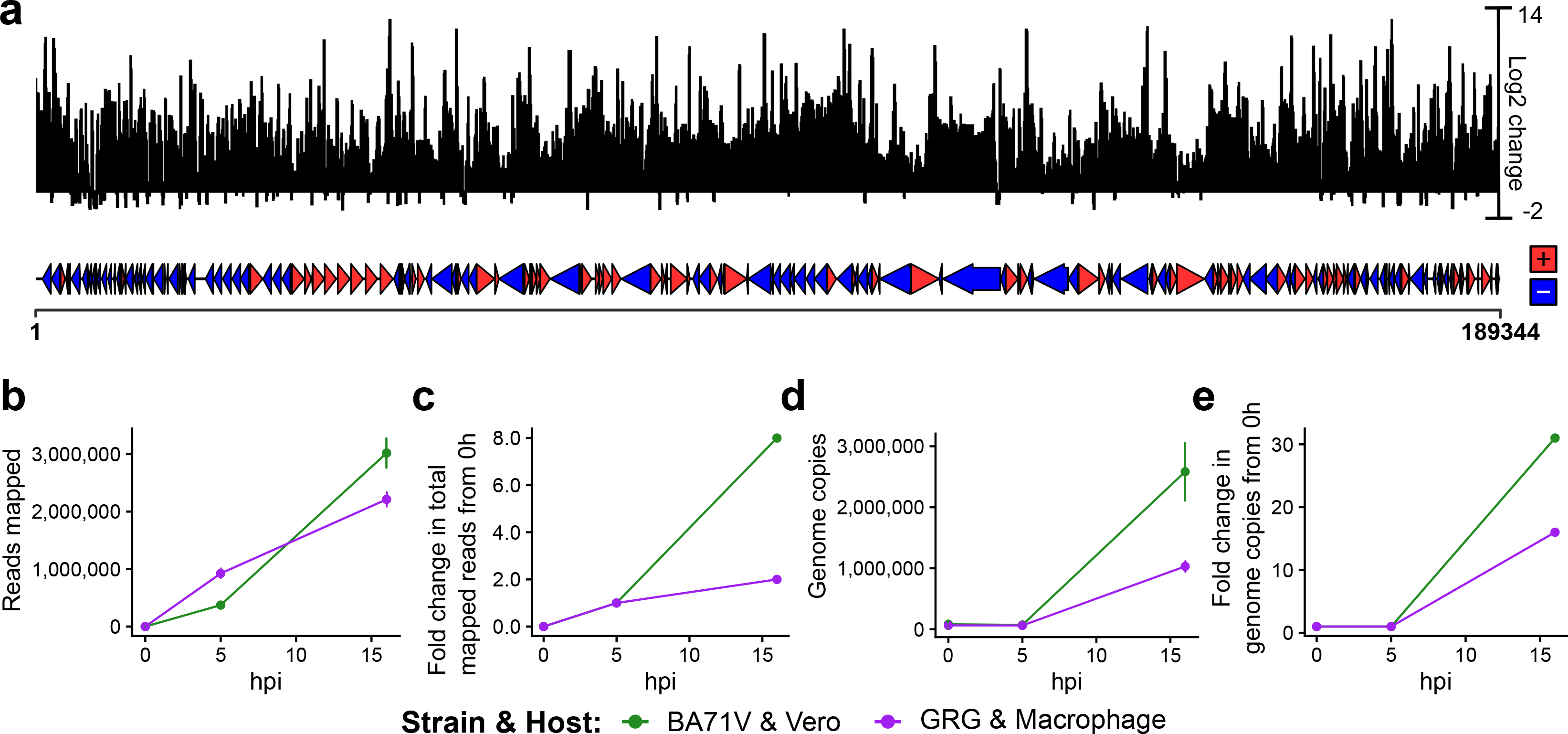
Increase in virus genome copy number mRNA levels during late infection. (a) The ‘log2 change’ represents log2 of the ratio of CAGE-seq reads (normalised per million mapped reads) at 16 hpi vs. 5 hpi per nucleotide across the genome. Alignment comparisons and calculations were done with deepTools (112). (b) Replicate means of CAGE-seq reads mapped to either the BA71V (green) or GRG (purple) genomes throughout infection. (c) Fold change in CAGE-seq reads during infection, calculated via mean value across 2 replicates, but with the assumption number of reads at 0 hpi is 0, therefore dividing by values from 5 hpi. (d) Change in genome copies from DNA qPCR of B646L gene, dividing by value at 0 hpi to represent ‘1 genome copy per infected cell’. (e) Fold change in genome copies present at 0 hpi, 5 hpi and 16 hpi from qPCR in (d). (d) calculated as for (c), but with actual vales for 0 hpi.

This observation can at least in part be attributed to the larger number of viral genomes during late infection, with increased levels of viral RNAP and associated factors available for transcription, following viral protein synthesis. Viral DNA-binding proteins, such as histone-like A104R (51), may remain associated with the genome originating from the virus particle in early infection. This could suppress spurious transcription initiation, compared to freshly replicated nascent genomes that are highly abundant in late infection. In order to test whether the increased mRNA levels correlated with the increased number of viral genomes in the cell, we determined the viral genome copy number by using quantitative PCR (qPCR against the p72 capsid gene sequence) using purified total DNA from infected cells isolated at 0 hpi, 5 hpi and 16 hpi, and normalized values to the total amount of input DNA. Using this approach, we observed genome copy levels that were consistent from 0 hpi to 5 hpi, consistent with this being pre-DNA replication, followed by a substantial increase at 16 hpi, which was more pronounced in BA71V infection (Figure 4d). This corresponded to a 15-fold increase in GRG genome copy numbers from late, compared to early times post-infection of porcine macrophages, and a 30-fold increase in BA71V during infection of Vero cells (Figure 4e). In summary, the ASFV transcriptome changes both qualitatively and quantitatively as infection progresses, and the increase of virus mRNAs during late infection is accompanied by the dramatic increase in viral genome copies. Interestingly, the increase in viral transcripts and genome copies was less dramatic in the virulent GRG strain.

### Correcting the bias of temporal expression pattern

The standard methods of defining differential gene expression are well established in transcriptomics using programs like DESeq2 (47). This is a very convenient and powerful tool which captures the nuances of differential expression in complex organisms. However, virus transcription is often characterised by more extreme changes, typically ranging from zero to millions of reads. Furthermore, in both BA71V and GRG strains the genome-wide mRNA levels and total ASFV reads increase over the infection time course (Figure 4 and Supplementary Table 1a). As a consequence, such normalisation against the total mapped transcripts per sample (RPM) generates overestimated relative expression values at 5 hpi, and understates those at 16 hpi (10). In order to validate the early-late expression patterns derived from CAGE-seq, we carried out RT-PCR for selected viral genes, as this signal is proportionate to the number of specific mRNAs regardless of the level of other transcripts – with the minor caveat that it can pick up readthrough transcripts from upstream genes. We tested differentially expressed conserved genes including GRG early- (MGF 505-7R, MGF 505-9R, NP419L), and D345L which showed stable relative expression values (RPM values in Figure 1e). All selected genes showed a consistently stronger RT-PCR signal during late infection in both BA71V and GRG (Figure 5a-d). The exception is NP419L whose levels were largely unchanged, and this is an example of how a gene whose transcript levels remain constant would be considered downregulated, when almost all other mRNA levels increase (Figure 5b).

**Figure 5.**
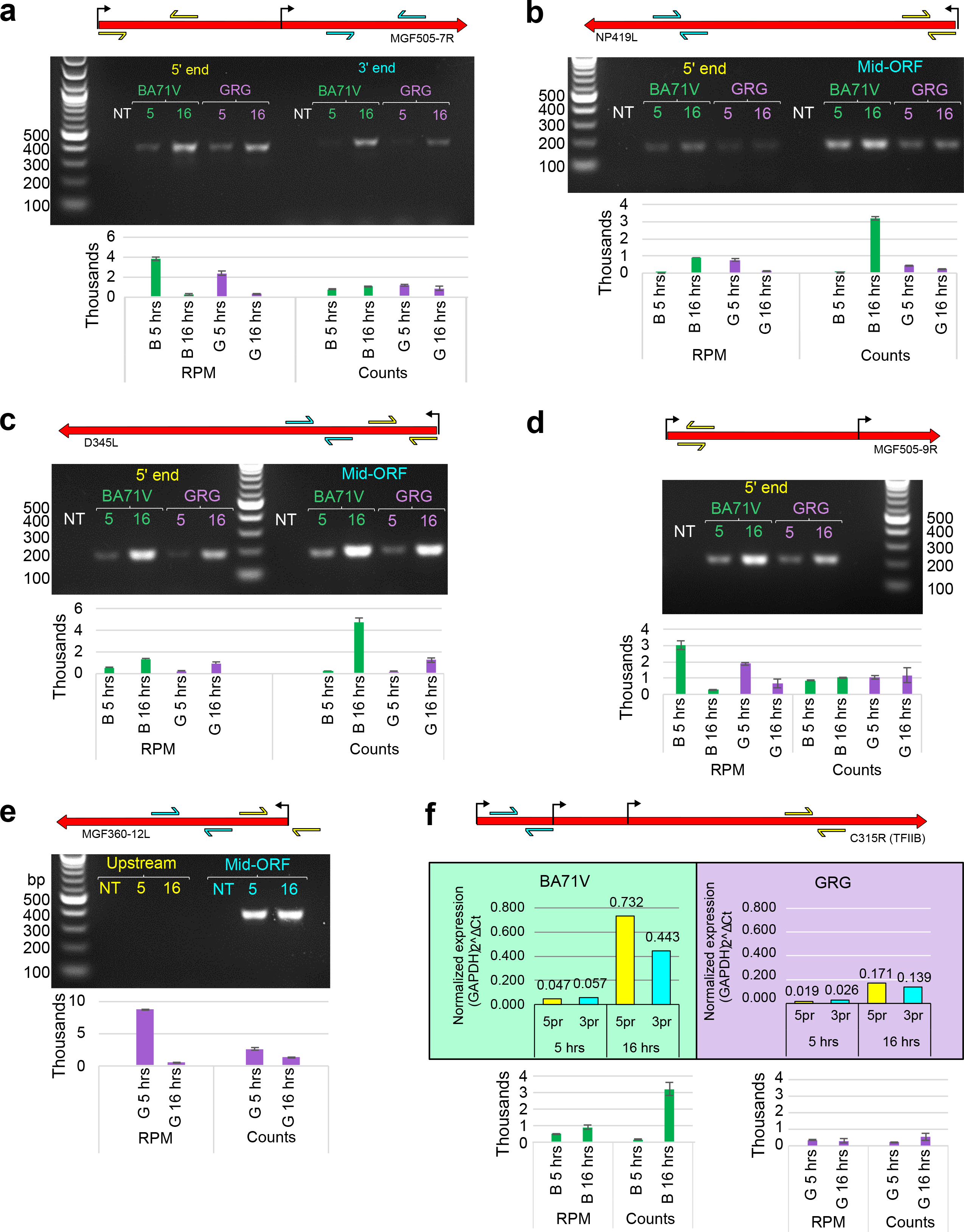
RT-PCR results of genes for comparison to CAGE-seq data from (a) MGF 505-7R, (b) NP419L, (c) D345L, (d) MGF 360-12L, (e) MGF 505-9R, and (f) qRT-PCR results of C315R (ASFV-TFIIB). (NT = no template control). For each panel at the top is a diagrammatic representation of each gene’s TSSs (bent arrow, including both pTSS and ioTSSs), annotated ORF (red arrow), and arrow pairs in cyan or yellow represent the primers used for PCR (see methods for primer sequences). Beneath each PCR results are bar charts representing the CAGE-seq results as either normalised (mean RPM) or raw (mean read counts) data, error bars show the range of values from each replicate.

The standard normalisation of NGS reads against total mapped reads (RPM) is regularly used as it enables a statistical comparison between samples and conditions, subject to experimental variations (52). Keeping this in mind, we used an additional method of analysing the ‘raw’ read counts to represent global ASFV transcript levels that are not skewed by the normalisation against total mapped reads. Figure 5 shows a side-by-side comparison of RT-PCR results, and the CAGE-seq data normalised (RPM) or expressed as raw counts, beneath each RT-PCR gel. Unlike CAGE-seq, RT-PCR will detect transcripts originating from read-through of transcripts initiated from upstream TSS including intra-ORF TSS (ioTSSs). To detect such ‘contamination’ we used multiple primer combinations in upstream and downstream segments of the gene (Figure 5c, cyan and yellow arrows) to capture and account for possible variations. Overall, our comparative analyses shows that the normalised data (RPM) of early genes such as MGF 505-7R and 9R indeed skews and overemphasises their early expression, while the raw counts are in better agreement with the mRNA levels detected by RT-PCR. In contrast, late genes such as NP419L and D345L would be categorised as late using all three quantification methods, in agreement with GRG CAGE-seq but not BA71V from Figure 3c. We validated the expression pattern of the early GRG-specific gene MGF 360-12L (Figure 5e). While the RPM values indicated a very strong decrease in mRNA levels from early to late time points, the decrease in raw counts was less pronounced and more congruent with the RT-PCR analysis, showing a specific signal with nearly equal intensity during early and late infection. Lastly, we used qRT-PCR to quantify C315R transcript levels, as this was close to the early vs late threshold, (a log2fold change of 0 in Figure 3c), which showed again that qRT-PCR better agreed with the raw counts.

### An improved temporal classification of ASFV genes

Based on the considerations above, we prepared a revised classification of temporal gene expression of the genes conserved between the two strains based on raw counts. The heatmap in Figure 6a shows the mRNA levels at early and late infection stages of BA71V and GRG strains (all in duplicates) with the genes clustered into five subcategories (1 to 5, Figure 6a) according to their early and late expression pattern, which are shown in Figure 6b. Genes that are expressed at high or intermediate levels during early infection but that also show high or intermediate mRNA levels during late infection are classified as ‘early’ genes belonging to cluster-1 (8 genes, levels: high to high, H-H), cluster-4 (33 genes, mid to mid, M-M) and cluster-5 (16 genes, low-mid to low-mid, LM-LM). Genes with low or undetectable mRNA levels during early infection, which increase to intermediate or high levels during late infection are classified as ‘late’ genes and belong to cluster-2 (15 genes, low to high, L-H) and cluster-3 (60 genes, low to mid, L-M), respectively. Overall, the clustered heatmap based on raw counts shows a similar but more emphasised pattern compared to the normalised (RPM) data (compare Figure 6 and Supplementary Figure 1). Calculating the percentage of reads per gene, which can be detected at 16 hpi compared to 5 hpi, reveals only a small number of genes have most ( ≥70%) of their reads originating during early infection: 30 genes in the GRG strain and 5 genes in the BA71V strain. For over half of the BA71V-GRG conserved genes, 90-100 % of reads can be detected during late infection (Figure 6c). For all GRG genes, this generates a significant difference between the raw counts per gene between time-points (Figure 6d).

**Figure 6.**
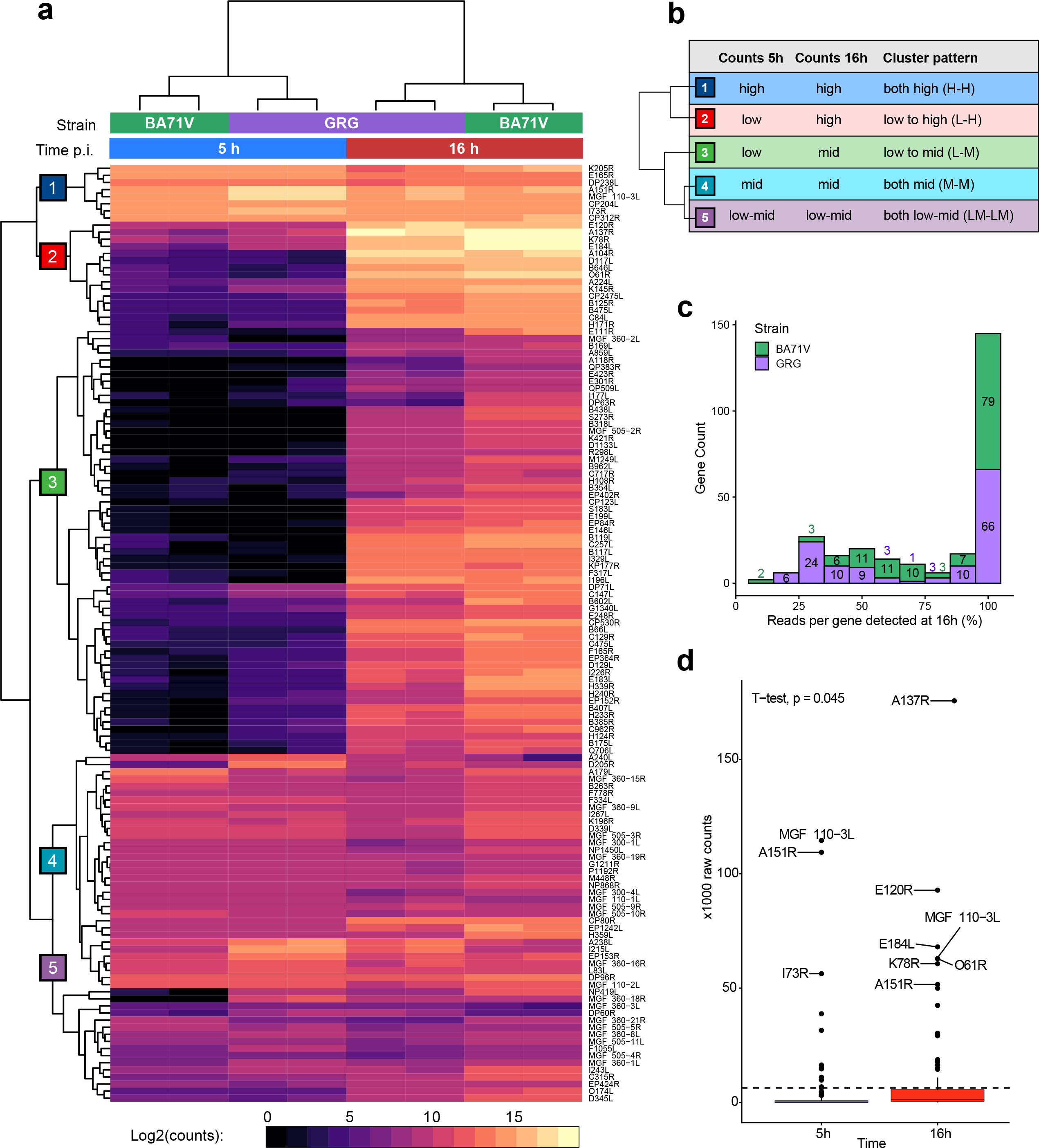
Comparison of the raw read counts for genes shared between BA71V and GRG. (a) clustered heatmap representation of raw counts for genes shared between BA71V and GRG, generated with pheatmap. (b) broad patterns represented by genes in the 5 clusters indicated in (a). (c) histogram showing the percentage of the total raw reads per gene which are detected at 16 hpi vs. 5 hpi post-infection, and comparing the distribution of percentages between GRG and BA71V. (d) Mean read counts from GRG at 5 hpi vs 16 hpi replicates, showing a significant increase (T-test, p-value: 0.045) from 5 hpi to 16 hpi.

Below we discuss specific examples of genes subcategorised in specific clusters. I73R is among the top twenty most-expressed genes during both early and late infection according to the normalised RPM values (Figure 2c and d) resides in cluster-1 (H-H) (Figure 6a). While I73R is expressed during early infection, the mRNA levels remain high with >1/3 of all reads detected during late infection in both strains when calculated as raw counts (34 % in GRG and 45 % in BA71V). This new analysis firmly locates I73R into cluster-1 (H-H) and is classified confidently as early gene. Notably, our new approach results in biologically meaningful subcategories of genes that are likely to be coregulated, e. g. the eight key genes that encode the ASFV transcription system including RNAP subunits RPB1 (NP1450L), RPB2 (EP1242L), RPB3 (H359L), RPB5 (D205R), RPB7 (D339L) and RPB10 (CP80R), the transcription initiation factor TBP (B263R) and the capping enzyme (NP868R) belong to cluster-4 (M-M), and transcription factors TFIIS (I243L) and TFIIB (C315R) belong to cluster-5 (LM-LM). The overall mRNA levels of cluster-4 and -5 genes are different, but remain largely unchanged during early and late infection, consistent with the transcription machinery being required throughout infection. In contrast, the mRNAs encoding the transcription initiation factors D6 (D1133L) and A7 (G1340L) are only present at low levels during early-but increase during late infection and thus belong to cluster-3 (L-M), classifying them as late genes. This is meaningful since the heterodimeric D6-A7 factor is packaged into viral particles (7), presumably during the late stage of the infection cycle. The mRNAs of the major capsid protein p72 (B646L) and the histone-like-protein A104R (51, 53) follow a similar late pattern but are present at even higher levels during late infection and therefore belong to cluster-2 (L-H).

### Architecture of ASFV promoter motifs

In order to characterise early promoter motifs (EPM) in the GRG strain, we extracted sequences 35 bp upstream of all early gene TSSs and carried out multiple sequence alignments. As expected, this region shows a conserved sequence signature in good agreement with our bioinformatics analyses of EPMs in the BA71V strain, including the correct distance between the EPM and the TSS (9-10 nt from the EPM 3’ end) and the ‘TA’ motif characteristic of the early gene Initiator (Inr) element (Figure 7a) (10). A motif search using MEME (54) identified a core (c)EPM motif with the sequence 5’-AAAATTGAAT-3’ (Figure 7b), within the longer EPM. The cEPM is highly conserved and is present in almost all promoters controlling genes belonging to cluster-1, -4 and -5 (Supplementary Table 3). A MEME analysis of sequences 35 bp upstream of late genes (Figure 7c), provided a 17-bp AT-rich core late promoter motif (cLPM, Figure 7d), however, this could only be detected in 46 of the late promoters.

**Figure 7.**
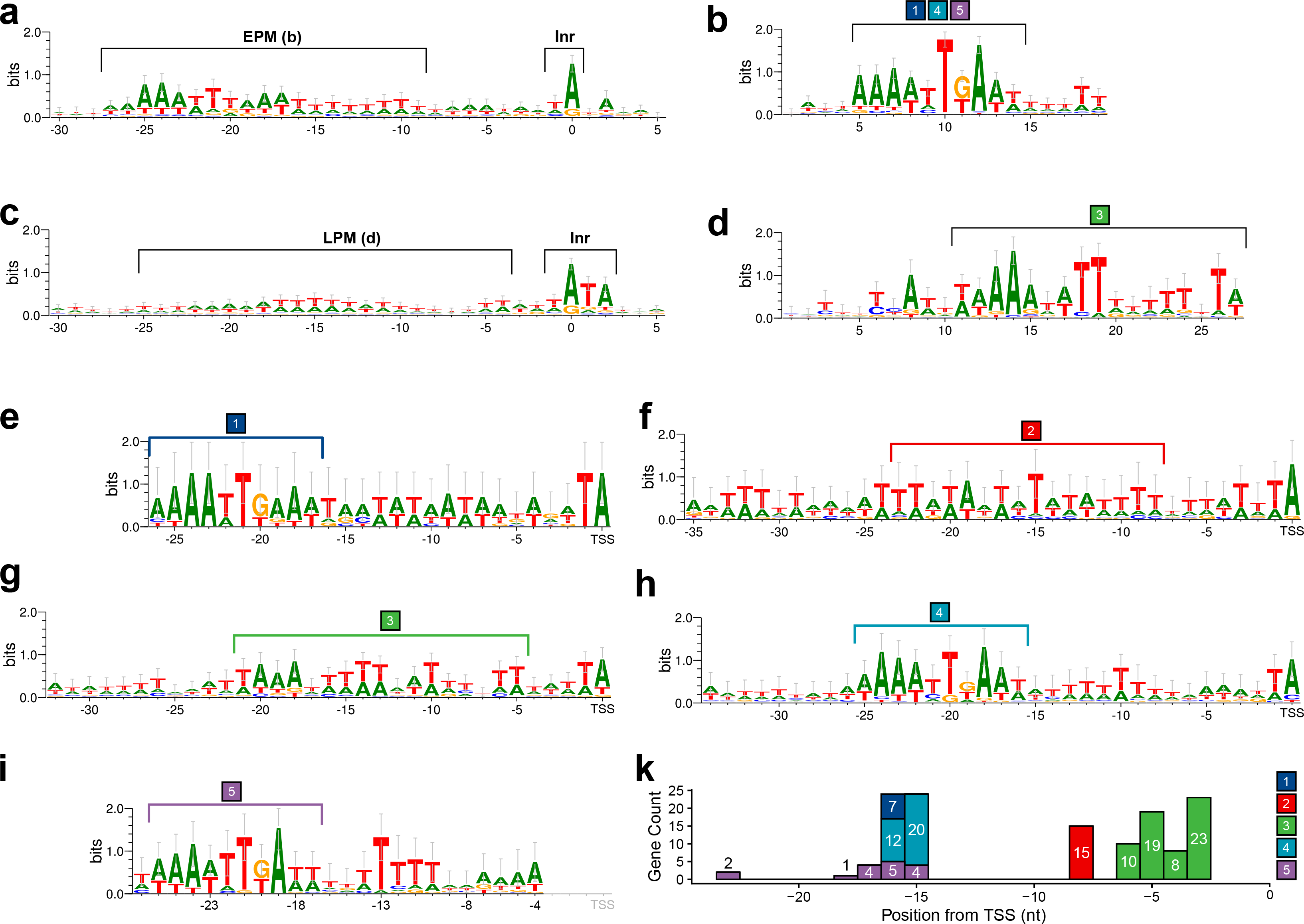
Promoter motifs and initiators detected in early and late ASFV GRG TSSs including alternative TSSs and those for nORFs. (a) Consensus of 30 bp upstream and 5 bp downstream of all 134 early TSSs including nORFs, with the conserved EPM (10) and Inr annotated. (b) 30 bp upstream and 5 bp downstream of all 234 late gene and nORFs TSSs, with the LPM and Inr annotated (c) The conserved EPM detected via MEME motif search of 35 bp upstream for 133 for 134 early TSSs (E-value: 3.1e-069). The conserved LPM detected via MEME motif search of 35 bp upstream for 46 for 234 late gene TSSs (E-value: 2.6e-003). The locations of the EPM shown in (b) and LPM shown in (d) are annotated with brackets in (a) and (b), respectively. Motifs detected via MEME search of 35 bp upstream of genes in clusters from Figure 6: cluster 1 (7 genes, E-value: 9.1e-012), 2 (15 genes, E-value: 2.6e-048), 3 (60 genes, E-value: 1.0e-167), 4 (32 genes, E-value: 4.7e-105), 5 (16 genes, E-value: 5.7e-036), are shown in e-i, respectively. For ease of comparison, (e), (g), (i) and (f), (h) are aligned at TSS position. All motifs were generated using Weblogo 3 (113). (k) shows the distribution of MEME motif-end distances, from last nt (in coloured bracket), to their respective downstream TSSs.

In an attempt to improve the promoter motif analyses and deconvolute putative sequence elements further, we probed the promoter sequence context of the five clusters (clusters 1-5 in Figure 7e-i, respectively) of temporally expressed genes with MEME (Supplementary Table 3). The early gene promoters of clusters-1 (H-H), -4 (M-M) and -5 (LM-LM) are each associated with different expression levels, and all of them contain the cEPM located 15-16 nt upstream of the TSS with two exceptions that are characterized by relatively low mRNA levels (Figure 7k). Interestingly, cluster-2 (L-H) promoters are characterized by a conserved motif with significant similarity to eukaryotic TATA-box promoter element that binds the TBP-containing TFIID transcription initiation factor (Figure 7f highlighted with red bracket, detected via Tomtom (55) analysis of the MEME motif output). Cluster-3 (L-M) promoters contain a long motif akin to the cLPM, derived from searching all late gene promoter sequences, and which is similar to the LPM identified in BA71V (Figure 7d and g, green bracket). All motifs described in the cluster analysis above could be detected with statistically significance (p-value < 0.05) via MEME, in every gene in each respective cluster with only two exceptions: MGF 110−3L from cluster-1, and MGF 360-19R from cluster-4, for the latter see details below.

### Updating Genome Annotations using Transcriptomics Data

TSS-annotation provides a useful tool for re-annotating predicted ORFs in genomes like ASFV (10) where many of the gene products have not been fully characterized and usually rely on prediction from genome sequence alone. We have provided the updated ORF map of the GRG genome in GFF format (Supplementary Table 1f). This analysis identified an MGF 360-19R ortholog (Figure 8), demonstrating how transcriptomics enhances automated annotation of ASFV genomes by predicting ORFs from TSSs. The MGF 360-19R was included in subsequent DESeq2 analysis showing it was not highly nor significantly differentially expressed (Supplementary Table 1e). Another important feature is the identification of intra-ORF TSSs (ioTSSs) within MGF 360-19R that potentially direct the synthesis of N-terminally truncated protein variants expressed either during early or late infection. The presence of EPM and LPM promoter motifs lends further credence to the ioTSSs (Figure 8). Similar truncation variants were previously reported for I243L and I226R (56) and in BA71V (10). In addition, we detected multiple TSSs within MGF 360-19R encoding very short putative novel ORFs (nORF) 5, 7 or 12 aa residues long; since these ioTSSs were present in both early and late infection they are not all likely to be due to pervasive transcription during late infection.

**Figure 8.**
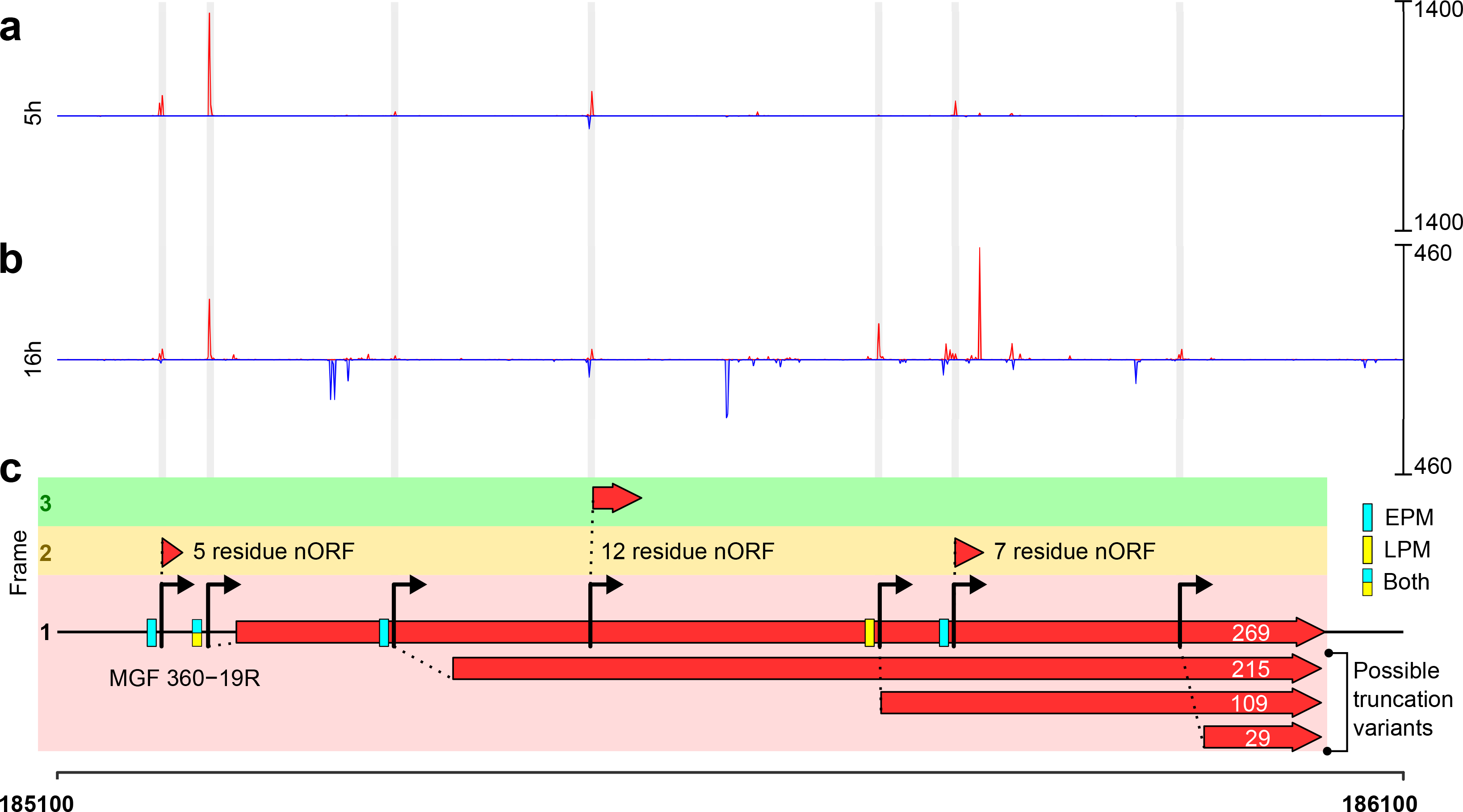
The TSSs of MGF 360-19R. Panels (a) 5 hpi and (b) 16 hpi show CAGE-seq 5’ end data from these time-points, in red are reads from the plus strand and blue from the minus strand, the RPM scales are on the right. (c) TSSs are annotated with arrows if they can generate a minimum of 5 residue-ORF downstream, and grey bars indicate where they are located on the CAGE-seq coverage in (a) and (b). ORFs identified downstream of TSSs are shown as red arrows (visualized with R package gggenes), including three short nORFs out of frame with MGF 360-19R. Also shown are three in-frame truncation variants, from TSSs detected inside the full-length MGF 360-19R 269-residue ORF, downstream of its pTSS at 185213. Blue or yellow boxes upstream of TSSs indicate whether the EPM or LPM (respectively) could be detected within 35 nt upstream of the TSS using FIMO searching (114).

We investigated the occurrence of ioTSS genome wide and uncovered many TSSs with ORFs downstream that were not annotated in the GRG genome (Supplementary Table 2a). These ORFs could be divided into sub-categories: in-frame truncation variants (Supplementary Table 2b, akin to MGF 360-19R in Figure 8), nORFs (Supplementary Table 2c), and simply mis-annotated ORFs. All updated annotations are found in Supplementary Table 1f. Putative truncation variants generated from ioTSSs were predominantly identified during late infection, suggesting these could be a by-product of pervasive transcription. Therefore, those detected early or throughout infection are perhaps more interesting, they span a variety of protein functional groups, and many gene-products are entirely uncharacterised (Figure 9a). The truncation variants additionally showed a size variation of 5’-UTRs between the ioTSSs and downstream start codon (Figure 9b). An example of a mis-annotation would be CP204L (Phosphoprotein p30, Figure 9c) gene that is predicted to be 201 residues long. The TSS determined by CAGE-seq and validated by Rapid Amplification of cDNA Ends (5’-RACE) is located downstream of the annotated start codon; based on our results we reannotated the start codon of CP204L which results in a shorter ORF of 193 amino acids (Figure 9c).

**Figure 9.**
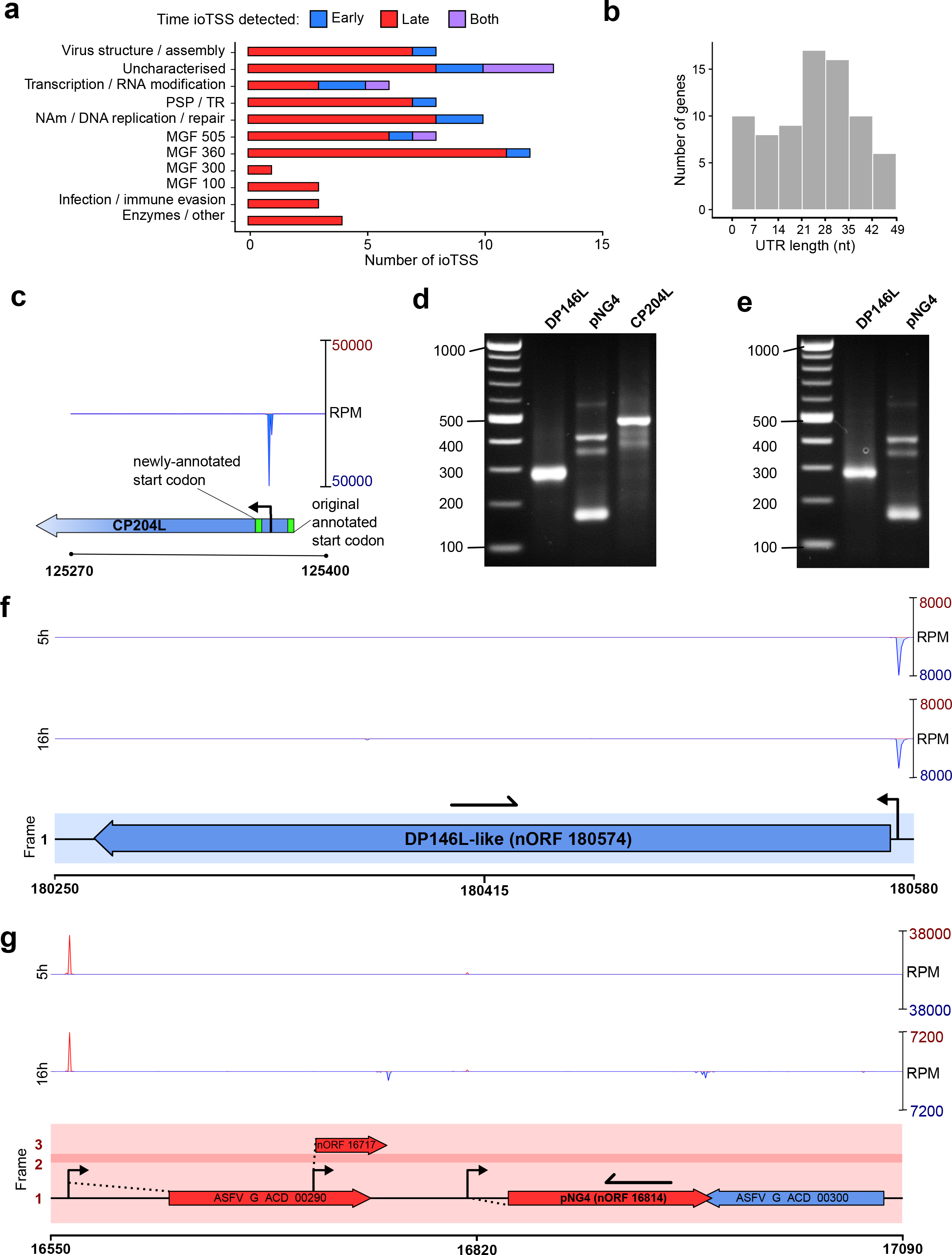
Summary of intra-ORF TSSs (ioTSSs) and nORFs detected in the GRG genome, further information in Supplementary Table 2. (a) Summarises the gene types in which ioTSSs were detected, showing an overrepresentation of MGFs, especially from families 360 and 505, furthermore, the majority of ioTSSs are detected at 16 hpi. (b) For ioTSSs in-frame with the original, summarised are the subsequent UTR lengths i.e. distance from TSS to next in frame ATG start codon, which could generate a truncation variant. (c) Example of a miss-annotation for CP204L, whereby the pTSS is downstream the predicted start codon. (d) and (e) show the results of 5’RACE for three genes (DP146L, pNG4, and CP204L, see methods for primers), at 5 hpi and 16 hpi, respectively. Examples of genome regions around DP146L (f) and pNG4 (g), wherein ioTSSs were detected with capacity for altering ORF length in subsequent transcripts, and therefore protein output. Primers used for 5’RACE for DP146L and pNG4 are represented as black arrows in (f) and (g), respectively.

Our GRG TSS map led to the discovery of many short nORFs, which are often overlooked in automated ORF annotations due to a minimum size, e. g. 60 residues in the original BA71V annotation (15). Some short ORFs have been predicted for the GRG genome including those labeled ‘ASFV_G_ACD’ in the Georgia 2007/1 genome annotation (19). However, their expression was not initially supported by experimental evidence, though we have now demonstrated their expression via CAGE-seq (Figure 2b, Supplementary Table 1e). We have now identified TSSs for most of these short ORFs, indicating at minimum they are transcribed. As described above, we noted that TSSs were found throughout the genome in intergenic regions in addition to those identified upstream of the 190 annotated GRG ORFs (including MGF 360-19R, Supplementary Table 2c). Our systematic, genome-wide approach identified 175 novel putative short ORFs. BLASTP (57) alignments showed that 13 were homologous to ORFs predicted in other strains, including DP146L and pNG4 from BA71V . We validated the TSSs for these candidates using 5’-RACE, which demonstrates the presence of these mRNAs and their associated TSSs at both time-points (Figure 9d and e, respectively), compared to our CAGE-seq data (Figure 9f and g, respectively).

### Putative single-SH2 domain protein encoding genes in MGF 100

Our understanding of the ASFV genome is hampered by the large number of genes with unknown functions. We approached this problem by searching for conserved domains of uncharacterised MGF members *in silico*. MGF 100 genes form the smallest multigene family and include three short (100– 150 aa) paralogs located at both genome ends (right, R and left, L): 1R, 1L (MGF_100-2L or DP141L in BA71V), and 3L (DP146L in BA71V). We predicted the two highly similar GRG ORFs I7L and I8L (51% sequence identity) to belong to the MGF 100 family (Figure 10a), as designated in the Malawi LIL20/1 strain (58). Both I7L and I8L show similar overall transcript levels to the annotated MGF 100 members -1L and 1R, though newly annotated MGF 100-3L (nORF_180573) was expressed at much higher levels. I7L and I8L are both early genes like MGF 100-3L, while MGF 100-1L and 1R are expressed late and not significantly changing, respectively (Supplementary Table 1e). Several lines of evidence suggest that I7L and I8L play an role during infection. I7L and I8L are expressed early and at high levels, their deletion along with L9R, L10L, and L11L ORFs reduces virulence in swine (59), and their loss is associated with the adaptation of the GRG2007/1 strain to tissue culture infection (60). To gain insight into the function of MGF family members including I7L and I8L, we generated computational homology models of MGF 100-1L -1R, I7L and I8L using Phyre2 (61) (Figure 10b). The structures selected by the algorithm for the modeling of MGF 100 proteins, included suppressor of cytokine signalling proteins 1 and -2, and the PI3-kinase subunit alpha, all of which are characterized by Src Homology 2 (SH2) domains (Figure 10b and Supplementary Table 2d). Canonical SH2 domains bind to phosphorylated Tyrosine residues and are an integral part of signalling cascades involved in the immune response (62). HHpred searches (63) predicted that indeed all MGF 100 members in BA71V and GRG include SH2 domains (Figure 10c).

**Figure 10.**
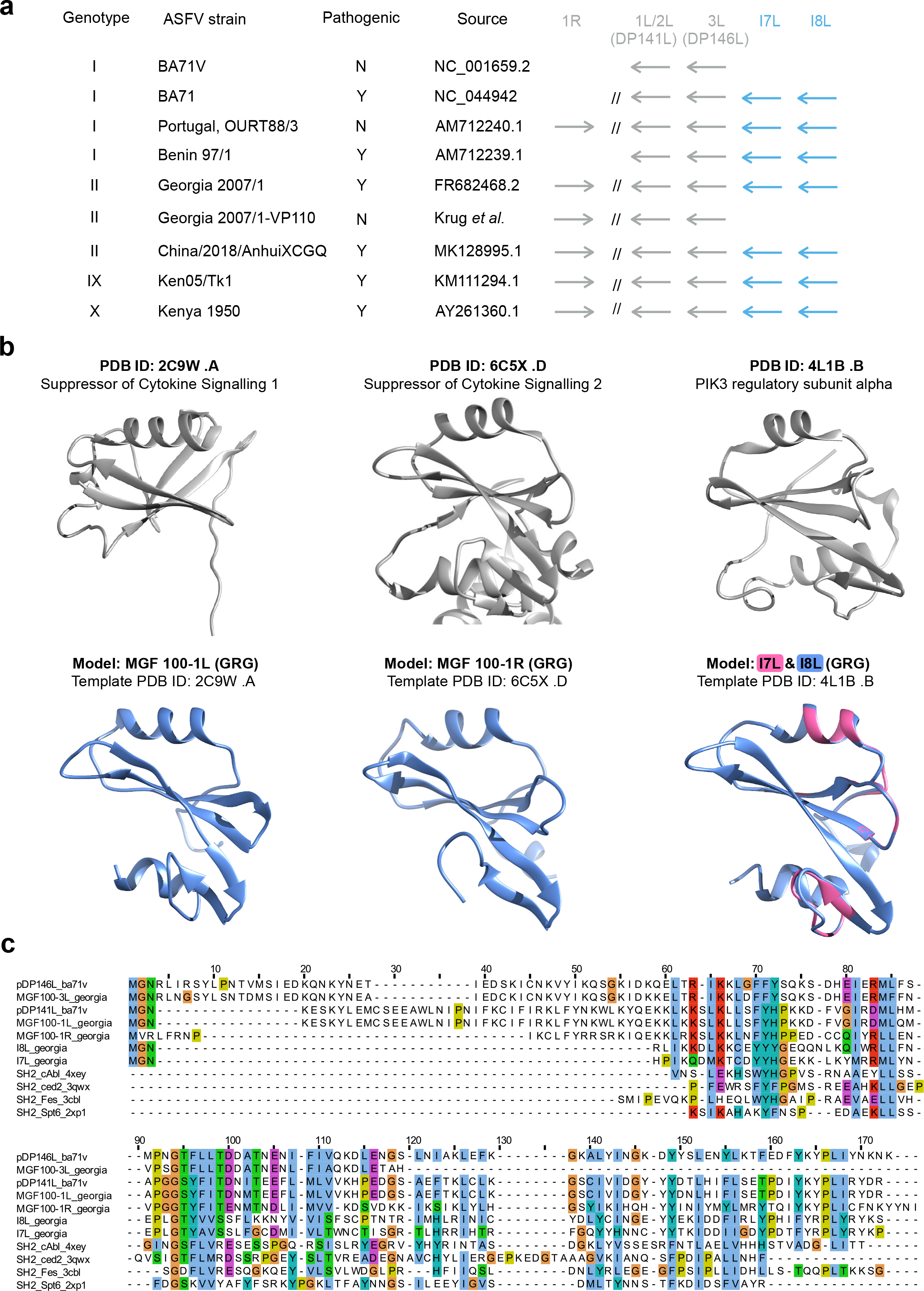
MGF 100 genes likely encode SH2-domain factors. (a) Occurrence of MGF 100 genes in selected ASFV strains, with genotype and pathogenicity indicated (as yes. Y, or no, N). ’1L/2L’ refers to the gene MGF 100-2L (DP141L in BA71V) and MGF 100-1L in the FR682468.1 genome annotation. (b) The top panel illustrates representative SH2 domain structures (Suppressor of Cytokine Signalling 1 and -2 and the PI3K alpha), and the bottom shows structural homology models of MGF 100 members 1L, 1R, and I7L and I8L superimposed. The PHYRE2 algorithm (56) was used to predict models for MGF 100 members (Supplementary Table 2d), and the structures at the top were detected as the top hits for each of the MGF 100 models shown in the lower panel. (c) Structure-guided multiple sequence alignment of selected MGF 100 member models, alongside known SH2 domain structures (annotated as SH2_name_PDB number).

### The response of the porcine macrophage transcriptome to ASFV infection

In order to evaluate the impact of ASFV on the gene expression of the host cell, we analysed transcriptomic changes of infected porcine macrophages using the CAGE-seq data from the control (uninfected cells), 5 hpi, and 16 hpi. We annotated 9,384 macrophage-expressed protein-coding genes with CAGE-defined TSSs (Supplementary Table 4). Although primary macrophages are known to vary largely in their transcription profile, the CAGE-seq reads were highly similar between RNA samples obtained from macrophages from two different animals in this study (Spearman’s correlation coefficients ≥ 0.77).

As TSSs are not well annotated for the swine genome, we annotated them *de novo* using our CAGE-seq data with the RECLU pipeline. 37,159 peaks could be identified, out of which around half (18, 575) matched unique CAGE-derived peaks annotated in Robert et al. (64) i.e. they were located closer than 100 nt to the previously described peaks. Mapping CAGE-seq peaks to annotated swine protein-coding genes led to identification of TSSs for 9,384 macrophage-expressed protein-coding genes (Supplementary Table 4). The remaining 11,904 swine protein-coding genes did not have assigned TSSs, and therefore their expression levels were not assessed. The majority of genes were assigned with multiple TSSs, and these TSS-assigned genes, corresponded to many critical functional macrophage markers, including genes encoding 56 cytokines and chemokines (including CXCL2, PPBP, CXCL8 and CXCL5 as the most highly expressed), ten S100 calcium binding proteins (S100A12, S100A8, and S100A9 in the top expressed genes), as well as interferon and TNF receptors (IFNGR1, IFNGR2, IFNAR1, IFNAR2, IFNLR1, TNFRSF10B, TNFRSF1B, TNFRSF1A, etc.), and typical M1/M2 marker genes such as TNF, ARG1, CCL24, and NOS2 (Supplementary Table 5

The 9,384 genes with annotated promoters were subjected to differential expression analysis using DESeq2 to compare the 5 and 16 hour infected cell time points with control non-infected cells (c, 5 and 16) in a pairwise manner i.e. between each condition. Expression of only 25 host genes was significantly deregulated between the control and 5 hpi, compared to 652 genes between 5 hpi and 16 hpi, and 1325 genes between mock-infected and 16 hpi (at FDR of 0.05). Based on the pairwise comparisons, we could distinguish major response profiles of the host genes. Late response genes, whose expression was significantly deregulated both between the uninfected control and 16 hpi and 5 and 16 hpi, and early response genes, whose expression was significantly deregulated between the control and 5 hpi, but not 5-16 hpi (Figure 11a). The latter category included only 20 genes, whereas more than 500 genes showed the late differentially regulated response: 344 genes were up-regulated, and 180 genes were down-regulated. The majority of the > 9000 genes analysed therefore were not differentially regulated. Comparison of differences between expression levels in the different samples indicate that macrophage differentially expressed transcription programs change mostly between 5 and 16 hpi (Figure 11b and c). The upregulated late response genes with highest expression levels included several S100 calcium binding proteins. In contrast, expression of important cytokines (including CCL24, CXCL2, CXCL5 and CXCL8) significantly decreased from 5 hpi to 16 hpi (Figure 11d).

**Figure 11.**
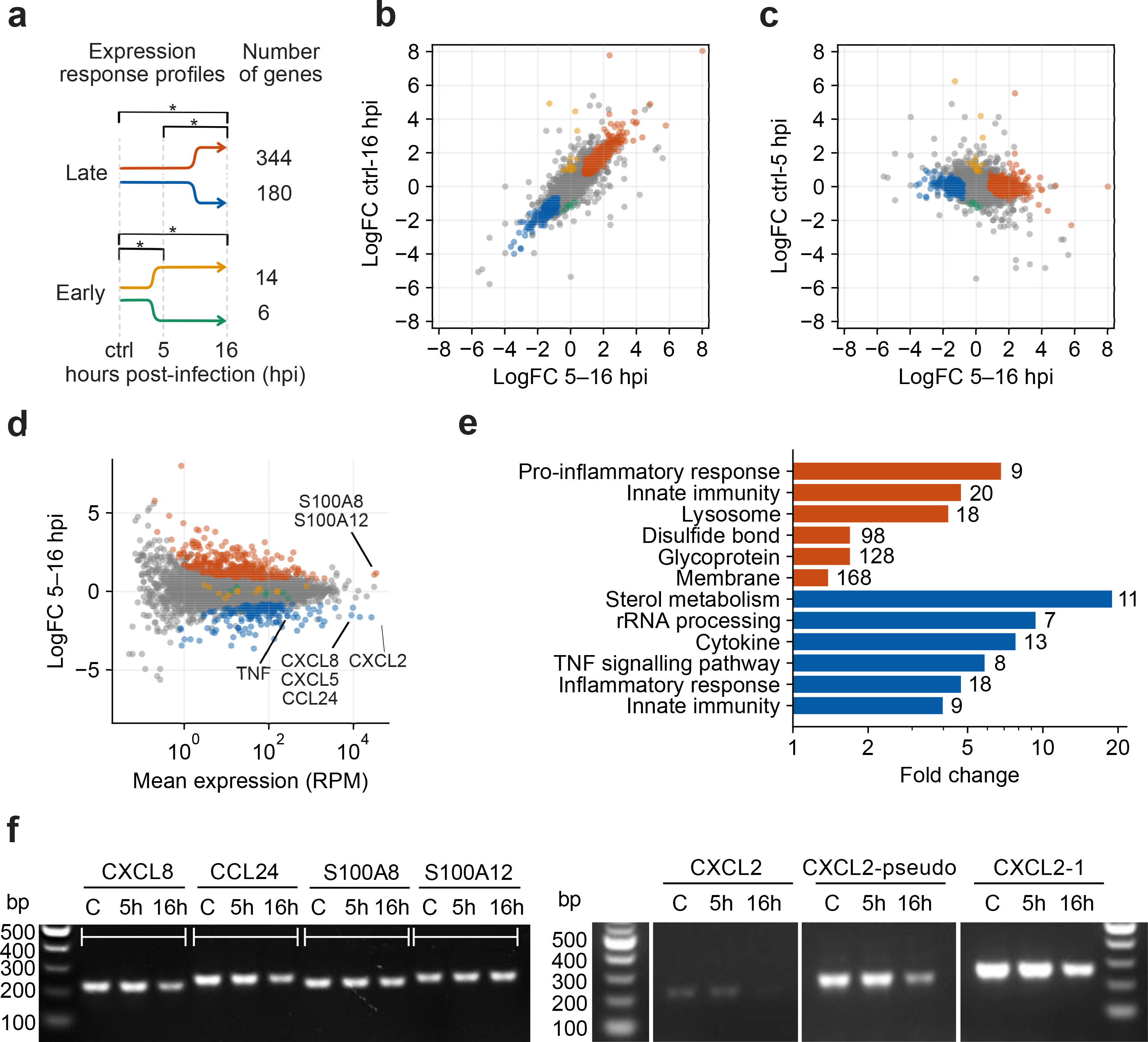
Changes in the swine macrophage transcriptome upon ASFV GRG infection. (a) Major expression response profiles of the pig macrophage transcriptome. Late response genes are significantly deregulated (false discovery rate < 0.05) in one direction both between mock-infected (ctrl) and 16 hpi as well as between 5 and 16 hpi, but not between mock-infected and 5 hpi. Early response genes are significantly deregulated in one direction both between ctrl and 5 hpi as well as ctrl and 16 hpi, but not between 5 and 16 hpi. (b) Relationship of log fold changes (logFC) of TSS-derived gene expression levels of the total 9,384 swine genes expressed in macrophages between 5– 16 hpi and ctrl–16 hpi. Colors correspond to the response groups from the panel a. (c) Relationship of log fold changes of TSS-derived gene expression levels of the total 9,384 swine genes expressed in macrophages between 5–16 hpi and ctrl–5 hpi. (d) MA plot of the TSS-derived gene expression levels between 5 and 16 hpi based on differential expression analysis with edgeR (108, 115). (e) Representative overrepresented functional annotations of the upregulated (red) and downregulated (blue) macrophage genes following late transcription response (Benjamini-corrected p-value lower than 0.05). Numbers on the right to the bars indicate total number of genes from a given group annotated with a given annotation. (f) RT-PCR of four genes of interest indicated in (d). ‘C’ is the uninfected macrophage control, NTC is the Non Template Control for each PCR, excluding template DNA. See methods for primers used.

To investigate the transcriptional response pathways and shed light on possible transcription factors involved in the macrophage response to ASFV infection, we searched for DNA motifs enriched in promoters of the four categories of deregulated genes in Figure 11a. Both late response promoter sets were significantly enriched with motifs, some of which contained sub-motifs known to be recognised by human transcription factors (Supplementary Figure 2). The highest-scored motif found in promoters of upregulated genes contained a sub-motif recognised by a family of human interferon regulatory factors (IRF9, IRF8 and IRF8, (Supplementary Figure 2a) that play essential roles in the anti-viral response. Interestingly, both upregulated and downregulated promoters (Supplementary Figure 2b and c, respectively) were enriched with extended RELA/p65 motifs. p65 is a Rel-like domain-containing subunit of the NF-kappa-B complex, regulated by I-kappa-B, whose analog is encoded by ASFV. This pathway being a known target for ASFV in controlling host transcription (65–68).

To understand functional changes in the macrophage transcriptome, we also performed gene set enrichment analysis using annotations of human homologs. The top enriched functional annotations in the upregulated late response genes include glycoproteins and disulfide bonds, transmembrane proteins, innate immunity, as well as positive regulation of inflammatory response (Figure 11e). In contrast, sterol metabolism, rRNA processing, cytokines, TNF signalling pathway, inflammatory response as well as innate immunity were the top enriched functional clusters among the downregulated late response genes. Interestingly, the genes associated with innate immunity appear overrepresented in both up-and downregulated gene subsets, yet cytokines are 8-fold enriched only in the downregulated genes). The mRNA levels of genes of interest were additionally verified using RT-PCR (Figure 11f).

### Protein expression of selected genes

In order to determine whether the regulation exerted by GRG on host transcription of immunomodulatory genes could also translate to protein levels, we selected representative proteins whose genes showed significant changes. ISG15 expression, part of the antiviral response genes of the type I IFN stimulation pathway, was measured with Western blot (Figure 12a), with ASFV infection being monitored via P30 levels (Figure 12b). Cytokines released from infected PAMS were quantified using ELISA tests for pig cytokines, TNF-α, CXCL8 and CCL2 (Figure 12c, d and e, respectively). As shown in Figure 12, the release/expression for all the tested proteins during GRG infection were similar or decreased in comparison to the control uninfected cells at both 5 hpi and 16 hpi, while the production of viral protein P30 increased, confirming an effective viral infection.

**Figure 12.**
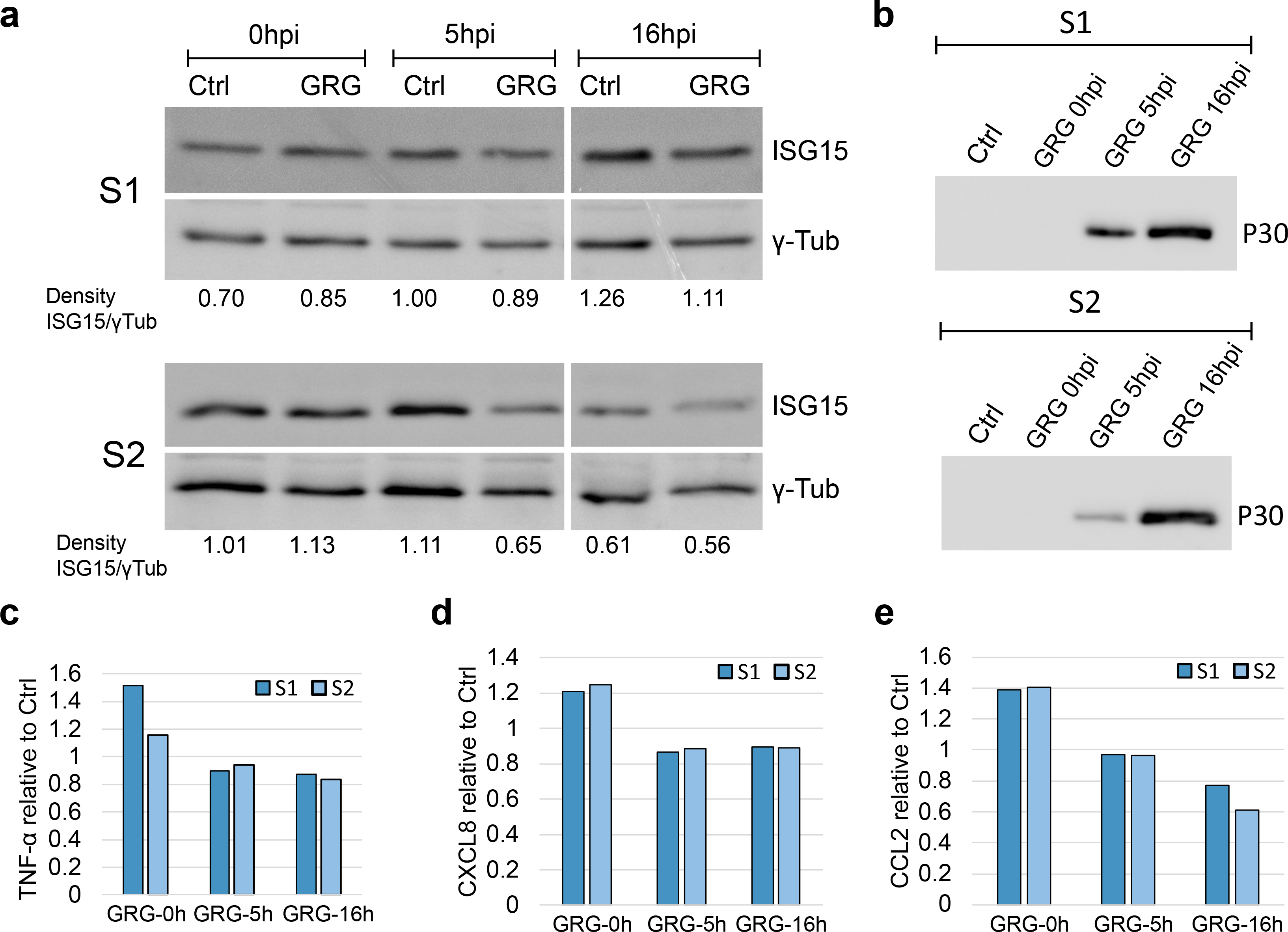
Protein expression at different times during infection of swine macrophages with ASFV-GRG. Two different batches of macrophages (S1 and S2) were infected with MOI 5 or left uninfected as a control (Ctrl) and at 0, 5 and 16hpi cellular extracts were collected and analysed via SDS-PAGE Western blot for the presence of ISG15 and γ-Tubulin as a protein loading control (a) and for the presence of viral protein P30 as control of ASFV infection (b). (c), (d) and (e) are the results from ELISAs for detection of porcine TNF-α, IL-8/CXCL8, and CCL2/MCP-1, respectively, in culture supernatants. Results are presented as ’Relative to control’ values (y-axis of c-e) calculated by performing ELISAs in parallel for control and GRG infection at each timepoint.

## Discussion

In order to shed light on the gene expression determinants for ASF virulence, we focussed our analyses on the similarities and differences in gene expression between a highly virulent Georgia 2007/1 isolate and a nonvirulent, lab-adapted strain BA71V. Previous annotation identified 125 ASFV ORFs that are conserved between all ASFV strain genomes irrespective of their virulence (16). They represent a ‘core’ set of genes required for the virus to produce infectious progeny and include gene products like those involved in virus genome replication, virion assembly, RNA transcription and modification. These genes are located in the central region of the genome (Figure 1a). Besides such essential genes, about one third are non-essential for replication, but have roles in evading host defence pathways. Some genes are conserved between isolates, but not necessarily essential core genes, for example apoptosis inhibitors: Bcl-2 family member A179L and IAP family member A224L (69). Other non-essential genes, especially MGF members, vary in number between isolates. Our transcriptomic analysis captured TSS signals from 119 genes both shared between the BA71V and GRG genomes, which also matched expression patterns during early and late infection, according to CAGE-seq (Figure 3, Figure 4a-c). Outliers include DP148R, which is unsurprising, given its promoter region is deleted in BA71V, and its coding region is interrupted by a frame shift mutation, therefore functional protein expression unlikely. DP148R is a non-essential, early-expressed virulence factor in the Benin 97/1 strain (49) – consistent with our GRG data. Many additional GRG genes, lost from BA71V are MGFs, which are mostly upregulated during early infection and located at the ends of the linear genome (Figure 1a). MGFs have evolved on the virus genome by gene duplication, and do not share significant similarity to other proteins, though some conserved domains, including ankyrin repeats, are present in some MGF 360 and 505 family members (17, 19).

Using advanced sequence searches and computational homology modelling we predict the members of the MGF 100 family to encode SH2 domains, including I7L and I8L. Although SH2 domains are primarily specific to eukaryotes, rare cases of horizontally transferred SH2 domains found in viruses, are implicated in hijacking host cell pTyr signalling (70). A large family of ‘super-binding’ SH2 domains were discovered in Legionella. Its members, including single SH2 domain-proteins are likely effector proteins during infection (71). We also identified a further MGF 100 member in the GRG genome as one of our nORFs, a partial 100-residue copy of DP146L (MGF 100-3L) (Supplementary Table 2c). Unlike its annotated MGF 100-1L and MGF 100-1R cousins it was downregulated from 5 hpi to 16 hpi (Supplementary Table 1e). Together with I7L and I8L, GRG encodes a total of 5 MGF 100 genes (Figure 10a). Interestingly, loss of MGF 100 members was observed during the process of adapting a virulent Georgia strain to grow in cultured cell lines (60). Deletion of MGF 100-1R, from a virulent genotype II Chinese strain (72) or of l8L from Georgia 2010 was shown not to reduce virulence of the virus in pigs or reduce virus replication in porcine macrophages (73). However, simultaneous deletion of genes l7L, l8L, l9L, l10L and l11L from a Chinese virulent isolate reduced virulence and surviving pigs were protected against challenge (59). In summary, although deletion of some individual MGF 100 genes does not lead to attenuation, deletion of l7L and l8L, in combination with l9L, l10L, and l11L did have an impact.

The Georgia 2007/1 genome was recently re-sequenced which identified a small number of genome changes affecting mapped ORFs and identified new ORFs (18). Adjacent to the covalently cross-linked genome termini, the BA71V genome contains terminal inverted repeats of >2 kbp, in which two short ORFs were identified (DP93R, DP86L). These were not included in previous GRG sequence annotations, however our nORFs included a 55-residue homolog of DP96R, which was a late, but not highly expressed gene. These are yet further examples of how transcriptomics aid in improving ASFV genome annotation. Functional data is available for only a few of proteins coded by ORFs not conserved between BA71V and GRG. This includes the p22 protein (KP177R), which is expressed on the cell membrane during early infection, and also incorporated into the virus particle inner envelope. The function of the KP177R-like GRG gene l10L has not been studied, but may provide an antigenically divergent variant of P22, enabling evasion of the host immune response (19). We found KP177R was highly expressed at 16 hpi, while l10L was also expressed late, but at much lower levels. Their function is unknown, though the presence of an SH2 domain indicates possible roles in signalling pathways (7,19,74).

MGF 110 members are among the highest expressed genes during early infection both in GRG (this study), and in BA71V (10), suggesting high importance during infection, at least in porcine macrophages and Vero cells, respectively. However, MGF 110 remains poorly characterised, and 13 orthologues were identified thus far, with numbers present varying between isolates (30). MGF 110 proteins possess cysteine-rich motifs, optimal for an oxidizing environment as found in the endoplasmic reticulum (ER) lumen or outside the cell, and MGF 110-4L (XP124L) contains a KDEL signal for retaining the protein in the ER (75). Since highly virulent isolates have few copies of these genes (for example, only 5 in the Benin 97/1 genome), it was assumed they are not importance for virulence in pigs (17), but their high expression warrant further investigation, which has recently begun in the form of deletion mutants. For example, deletion of MGF 110-9L from a Chinese genotype II virulent strain, reduced virulence (35), whereas deletion of MGF 110-1L from Georgia 2010 (76) did not substantially affect virulence.

There is however, good evidence that MGF 360 and 505 carry out important roles in evading the host type I interferon (IFN) response - the main host antiviral defence pathway (37). Evidence for the role of MGF 360 and 505 genes in virulence was obtained from deletions in tissue-culture adapted and field attenuated isolates, as well as targeted gene deletions This correlated with induction of the type I interferon response, which itself is inhibited in macrophages infected with virulent ASFV isolates (32,38,39). Deletions of these MGF 360, and 505 genes also correlated with an increased sensitivity of ASFV replication, to pre-treatment of the macrophage cells with type I IFN (40). Thus, the MGF 360 and 505 genes have roles in inhibiting type I IFN induction and increasing sensitivity to type I IFN. However, it remains unknown if these MGF 360 and MGF 505 genes act synergistically or if some have a more important role than others type I IFN suppression. Our DESeq2 analysis did show that members of both these families showed very similar patterns of early expression (Figure 2 and Figure 3), conserved cEPM-containing promoters, and almost exclusive presence in clusters-1 (H-H), -4 (M-M), and -5 (LM-LM) (Figure 6 and Figure 7), consistent with ASFV prioritising inhibition of the host immune response during early infection.

An interesting pattern which emerged during our CAGE-seq analysis was the clear prevalence of ioTSSs within ORFs, especially in MGFs (Figure 8 and Figure 9). However, it is not clear whether subsequent in-frame truncation variants generate stable proteins, nor what their function could be. Perhaps even more interesting was the discovery of 176 nORFs (including MGF 360-19R), with clear TSSs according to CAGE-seq, highlighting the power of transcriptomics to better annotate sequenced genomes. We were able to detect previously unannotated genes from other strains, and partial duplications of genes already encoded in GRG (Supplementary Table 2).

The increase in transcription across the ASFV genome during late infection (10), appears ubiquitous. At least 50 genes have previously been investigated in single gene expression studies using Northern blot or primer extension (for review see references (10, 77). Transcripts from over two thirds of these genes were detected during late infection, and a quarter had transcripts detected during both early and late infection. Therefore, clear evidence using several techniques now support this increase in ASFV transcripts at late times post-infection. It is not entirely clear whether it is due to pervasive transcription, high mRNA stability or a combination of factors. However, there is a correlated increase in viral genome copies, potentially available as templates for pervasive transcription. The increase in genome copies is more pronounced in BA71V compared to GRG, which likewise is reflected in the increase in transcripts during late infection (Figure 4).

Our transcriptomic analysis of the porcine macrophage host revealed 522 genes whose expression patterns significantly changed between 5 and 16 hrs post-infection (Figure 11a) and only 20 genes were found to change between the control cells and those infected for 5 hpi. In aggregate, this reflects a relatively slow host response to ASFV infection following expression of early ASFV genes. We observed mild downregulation of some genes e.g. ACTB coding for ß-actin, eIF4A, and eIF4E (Supplementary Table 5), resembling patterns previously shown by RT-qPCR (78). The macrophage transcriptome mainly shuts down immunomodulation between 5 hpi to 16 hpi post-infection; cytokines appeared highly expressed at 5 hpi, but downregulated from 5 hpi to 16 hpi. Of the 54 cytokine genes we detected, expression of thirteen was decreased: four interleukin genes (IL1A, IL1B, IL19, IL27), four pro-inflammatory chemokines (CCL24, CXCL2, CXCL5, CXCL8), and tumor necrosis factor (TNF) genes. Since inflammatory responses serve as the first line of host defense against viral infections, viruses have developed ways to neutralise host pro-inflammatory pathways. ASFV encodes a structural analog of IκB, A238L, which was proposed to act as a molecular off-switch for NFκB-targeted pro-inflammatory cytokines (67). In our study, A238L is one of the most expressed ASFV genes at 5 hpi, but significantly downregulated afterwards (Figure 2c). Accordingly, swine homologs of human NFκB target genes were significantly over-represented (3.8 fold) among downregulated macrophage genes (Fisher’s exact p-value < 1e-5, based on human NFκB target genes from https://www.bu.edu/nf-kb/gene-resources/target-genes/). Downregulated genes include interleukins 1A, 1B, and 8, and 27 (IL1A, IL1B, CXCL8, IL27), TNF, as well as a target for common nonsteroidal anti-inflammatory drugs, prostaglandin-endoperoxide synthase 2 (PTGS2 or COX-2) (Supplementary Figure 2). Interestingly, promoters of both up-and downregulated genes contained a motif with the sequence preferentially recognised by the human p65-NFκB complex (79). Expression of TNF, a well-known marker gene for acute immune reaction and M1 polarisation, was recorded at a high level in control samples and at 5 hpi, but significantly dropped at 16 hpi. It has been already shown that ASFV inhibits transcription of TNF and other proinflammatory cytokines (67). On the other hand, the downregulation of TNF stands in contrast to previous results from ASFV-E75 strain-infected macrophages in vitro, where TNF expression increased significantly after 6 hpi (80). Therefore, the different time courses of TNF expression induced by the moderately virulent E75 and more virulent Georgia strain may reflect different macrophage activation programs (81).

We investigated if the modulation of transcription we observed by CAGE-seq during GRG infection of PAMS was also observed at the protein level. We analysed the secretion or expression of different immunomediators (cytokines CCL2, CXCL8, TNF-α and interferon stimulated gene ISG15) at different times following infection of PAMS. We confirmed that that the infection did not lead to an increase of these mediators at either 5h or 16h infection. Secretion or expression of these proteins were similar or slightly decreased in infected cells in comparison to control non-infected cells. The results indicated that the control by virulent Georgia 2007/1 of host cell responses to infection we observed at the transcription level can lead to a control also at the level of the protein production. Interestingly, CCL2 transcription was somewhat upregulated at late infection (Supplementary Table 5), whereas its protein release to the supernatant was decreased (Figure 12e). ASFV has been shown to prioritize expression of its encoded proteins by sequestering components of the host translation machinery to viral factories (82). The levels or functions of host proteins may also be modulated by targeting for post-translational modification or degradation (82–84). Therefore, in addition to control at the transcriptional level ASFV may modulate the production of immunomodulatory host proteins at a later step, as seems to occur for CCL2, a known chemoattractant for myeloid and lymphoid cells (85), that could be an important target for regulation by ASFV.

Four S100 family members are among the host genes that are upregulated after 5 hpi (Figure 11b) including S100A8, S100A11, S100A12, and S100A13. S100A8 and S100A12 are among the most highly expressed genes on average throughout infection. S100 proteins are calcium-binding cytosolic proteins that are released and serve as a danger signal, and stimulate inflammation (86). Once released from the cell, S100A12 and S100A8 function as endogenous agonists to bind TLR4 and induce apoptosis and autophagy in various cell types (86). S100A8 and S100A9 were also found in the RNA-seq whole blood study as the top upregulated upon infection of the pigs with Georgia 2007/1, but not of a low pathogenic ASFV isolate OURT 88/3 (43).

Previous studies described global swine transcriptome changes upon ASFV infection using short read sequencing (Illumina): including the RNA-seq described above (43) and a microarray study of primary swine macrophage cell cultures infected with the GRG strain, at six time points post-infection (42). Although these varied in designs and selected methods, results of these works both give some indication into the main host immune responses and ways how ASFV could evade them. The latter microarray study indicated similar suppression of inflammatory response after 16 hpi as we observed in this study, with expression of many cytokines down-regulated relative to non-infected macrophages (42). More-recently, there have been several transcriptomic studies using classical RNA-seq of ASFV infections from Chinese isolates (44–46). Fan et al (44) investigated the transcriptomic and proteomic response within tissues of pigs following ASFV infection and death, though this was not directly comparable to our own analysis in PAMs, due to their observations being of a far later infection stage (post-mortem) than our 16 hrs time-point. The two most-comparable studies to ours were carried out on a Chinese genotype II pathogenic strain during infection of PAMs. Ju et al. (45) investigated 6, 12 and 24 hpi, while Yang et al. (46) investigated 12, 24, and 36 hpi. However, comparison of the overlapping time points of 12 hpi and 24 hpi did not yield similar host gene expression changes, possibly due to variation among primary macrophages or due to the low MOI of 1 used in both studies. In summary, these differences highlight that our understanding of the host-virus relationship during ASFV infection is still not well understood, and further work is needed to understand why such substantial variation in host gene expression can arise.

A further important note, is that all of the studies described above are using classical RNA-seq-based methods, the nucleotide resolution of which, is not sufficient to investigate differential expression of both the virus and host simultaneously. Investigating the viral transcriptome is especially difficult in a compact genome like that of ASFV, where transcription read-through can undermine results from classical RNA-sequencing techniques (10, 87). A recent investigation into ASFV RNA transcripts using long-read based Oxford Nanopore Technologies (ONT) – provides fascinating insight into their length and read-through heterogeneity. This new method highlighted how misleading short read sequencing with classical RNA-seq can be when quantifying ASFV gene expression, due to the abundance of readthrough occurring in ASFV, generating transcripts covering multiple viral ORFs. This study did however, unfortunately lack the read coverage for in-depth analysis of host transcripts alongside that of viral transcripts (88, 89).

Here we have demonstrated that CAGE-seq is an exceptionally powerful tool for quantifying relative expression of viral genes across the ASFV genome, as well as making direct comparison between strains for expression of shared genes, and further highlighting the importance of highly-expressed but still functionally uncharacterised viral genes. CAGE-seq conveniently circumvents the issue in compact viral genomes like those of ASFV and VACV, of transcripts reading through into downstream genes which cannot be distinguished from classical short-read RNA-seq (10,43,90). Furthermore, it enables us to effectively annotate genome-wide, the 5’ ends of capped viral transcripts, and thus TSSs of viral genes, and subsequently their temporal promoters. This 5’ end resolution in ASFV is still not achievable via ONT long read sequencing (88, 89). We have now expanded on promoter motifs we previously described (Figure 7), to identify 5 clusters of genes (Figure 6), with distinct patterns of expression. Three of these clusters (-1: high to high levels, -4: mid to mid, and -5 low-mid to low-mid) have slightly differing promoters, with a highly conserved core EPM. This is akin to the early gene promoter of VACV (87) for VETF recognition and early gene transcription initiation (13,91,92). We have found late genes can be categorised into two types that either increase from low to extremely high expression levels (e. g. p72-encoding B646L) in cluster-2, or from low to medium expression levels in cluster 3 (e. g VETF-encoding genes). The promoters of these genes show resemblance to the eukaryotic TATA-box (93) or the BA71V LPM (10), respectively. Our analysis additionally shows the potential for a variety of non-pTSSs: alternative ones used for different times in infection, ioTSSs which could generate in-frame truncation variants of ORFs, sense or antisense transcripts relative to annotated ORFs, and finally TSSs generating nORFs, which predominantly have no known homologs.

In summary, it is becoming increasingly clear that the transcriptomic landscape of ASFV and its host during infection is far more complex than originally anticipated. Much of this raises further questions about the basal mechanisms underlying ASFV transcription and how it is regulated over the infection time course. Which subsets of initiation factors enable the RNAPs to recognise early and late promoters? Does ASFV include intermediate genes, and what factors enables their expression? What is the molecular basis of the pervasive transcription during late infection? The field of ASFV transcription has been understudied and underappreciated and considering the severe threat that ASF poses for the global food system and -food security, we now need to step up and focus our attention and resources to study the fundamental biology of ASFV to develop effective antiviral drugs and vaccines.

## Methodology

### GRG-Infection of Macrophages and RNA-extraction

Primary porcine alveolar macrophage cells were collected from two animals following approval by the local Animal Welfare and Ethical Review Board at The Pirbright Institute. Cells were seeded in 6-well plates (2x10^6^ cells/well) with RPMI medium (with GlutaMAX), supplemented with 10% Pig serum and 100 IU/ml penicillin, 100 μg/ml streptomycin. They were infected as 2 replicate wells for 5 hpi or 16 hpi with a multiplicity of infection (MOI) of 5 of the ASFV Georgia 2007/1 strain, while uninfected cells were seeded in parallel as a control (mock-infection). Total RNA was extracted according to manufacturer’s instructions for extraction with Trizol Lysis Reagent (Thermo Fisher Scientific and the subsequent RNAs were resuspended in 50µl RNase-free water and DNase-treated (Turbo DNAfree kit, Invitrogen). RNA quality was assessed via Bioanalyzer (Agilent 2100). 5 μg of each sample was ethanol precipitated before sending to CAGE-seq (Kabushiki Kaisha DNAFORM, Japan). Samples were named as follows: uninfected cells or ’mock’ (C1-ctrl and C2-ctrl), at 5 hpi post-infection (samples G1-5h and G2-5h), and at 16 hpi post-infection (G3-16h and G4-16h).

### CAGE-sequencing and Mapping to GRG and *Sus scrofa* Genomes

Library preparation and CAGE-sequencing of RNA samples was carried out by CAGE-seq (Kabushiki Kaisha DNAFORM, Japan). Library preparation produced single-end indexed cDNA libraries for sequencing: in brief, this included reverse transcription with random primers, oxidation and biotinylation of 5’ mRNA cap, followed by RNase ONE treatment removing RNA not protected in a cDNA-RNA hybrid. Two rounds of cap-trapping using Streptavidin beads, washed away uncapped RNA-cDNA hybrids. Next, RNase ONE and RNase H treatment degraded any remaining RNA, and cDNA strands were subsequently released from the Streptavidin beads and quality assessed via Bioanalyzer. Single strand index linker and 3’ linker was ligated to released cDNA strands, and primer containing Illumina Sequencer Priming site was used for second strand synthesis. Samples were sequenced using the Illumina NextSeq 500 platform producing 76 bp reads. FastQC (94) analysis was carried out on all FASTQ files at Kabushiki Kaisha DNAFORM and CAGE-seq reads showed consistent read quality across their read-length, therefore, were mapped in their entirety to the GRG genome (FR682468.1) in our work using Bowtie2 (95), and *Sus scrofa* (GCF_000003025.6) genome with HISAT2 (95, 96) by Kabushiki Kaisha DNAFORM.

### Transcription Start Site-mapping Across Viral GRG Genome

CAGE-seq mapped sample BAM files were converted to BigWig (BW) format with BEDtools (97) genomecov, to produce per-strand BW files of 5’ read ends. Stranded BW files were input for TSS-prediction in RStudio (98) with Bioconductor (99) package CAGEfightR (100). Genomic feature locations were imported as a TxDb object from FR682468.1 genome gene feature file (GFF3). CAGEfightR was used to quantify the CAGE reads mapping at base pair resolution to the GRG genome - at CAGE TSSs, separately for the 5 hpi and 16 hpi replicates. TSS values were normalized by tags-per-million for each sample, pooled, and only TSSs supported by presence in both replicates were kept. TSSs were assigned to clusters, if within 25 bp of one another, filtering out pooled, RPM-normalized TSS counts below 25 bp for 5 hpi samples, or 50 bp for 16 hpi, and assigned a ‘thick’ value as the highest TSS peak within that cluster. A higher cut-off for 16 hpi was used to minimise the extra noise of pervasive transcription observed during late infection (10). TSS clusters were assigned to annotated FR682468.1 ORFs using BEDtools intersect, if its highest point (‘thick’ region) was located within 500 bp upstream of an ORF, ‘CDS’ if within the ORF, ‘NA’ if no annotated ORF was within these regions. Multiple TSSs located within 500 bp of ORFs were split into subsets: ‘Primary’ cluster subset contained either the highest scoring CAGEfightR cluster or the highest scoring manually-annotated peak (when manual ORF corrections necessary), and the highest peak coordinate was defined as the primary TSS (pTSS) for an ORF. Further clusters associated with these ORFs were classified as ‘non-primary’, with their highest peak as a non-primary TSS (npTSS). If the strongest TSS location was intra-ORF, without any TSSs located upstream of the ORF, then the ORF was manually re-defined as starting from the next ATG downstream.

### DESeq2 Differential Expression Analysis of GRG Genes

For analysing differential expression with the CAGE-seq dataset, a GFF was created with BEDtools extending from the pTSS coordinate, 25 bp upstream and 75 bp downstream, however, in cases of alternating pTSSs this region was defined as 25 bp upstream of the most upstream pTSS and 75 bp downstream of the most downstream pTSS. HTSeq-count (101) was used to count reads mapping to genomic regions described above for both the RNA-and CAGE-seq sample datasets. The raw read counts were then used to analyse differential expression across these regions between the time-points using DESeq2 (default normalisation described by Love et al. (47)) and those regions showing changes with an adjusted p-value (padj) of <0.05 were considered significant. A caveat of this ‘early’ or ‘late’ definition is that it is a binary definition of whether a gene is up-or downregulated between conditions (time-points), relative to the background read depth of reads, which map to the genome in question. Further analysis of ASFV genes used their characterised or predicted functions, from the VOCS tool database (https://4virology.net/) (102, 103) entries for the GRG genome.

### Quantification of viral genome copies at different time points of infection

Porcine lung macrophages were seeded and infected as described above. *Vero* cells were similarly cultured in 6-well plates in DMEM medium supplemented with 10% Fetal calf serum, 100 IU/ml penicillin and 100 μg/ml streptomycin, when semi-confluent they were infected with MOI 5 of Ba71V. Immediately after infection (after 1h adsorption period, considered ‘0 hpi), or at 5 hpi, and 16 hpi, the supernatant was removed and nucleic acids were extracted using the Qiamp viral RNA kit (Qiagen) and quantified using a NanoDrop spectrophotometer (ThermoFisher Scientific). For quantification of viral genome copy equivalents, 50 ng of each nucleic acid sample was used in qPCR with primers and probe targeting the viral capsid gene B646L. As previously described (104), standard curve quantification qPCR was carried out on a Mx3005P system (Agilent Technologies) using the primers CTGCTCATGGTATCAATCTTATCGA and GATACCACAAGATC(AG)GCCGT and probe 5′-(6-carboxyfluorescein [FAM])-CCACGGGAGGAATACCAACCCAGTG-3′-(6-carboxytetramethylrhodamine [TAMRA]).

### Analysis of mRNA levels by RT-PCR and quantitative real time PCR (qPCR)

RNA from GRG or Ba71V infected macrophages, or *Vero* cells respectively, or from uninfected cell controls, was collected at the different time points post-infection with Trizol, as described above. RNA was reverse transcribed (800 ng RNA per sample) using SuperScript III First-Strand Synthesis System for RT-PCR and random hexamers (Invitrogen). For PCR, cDNAs were diluted 1:20 with nuclease free water and 1 μl each sample was amplified in a total volume of 20 μl using Platinum™ Green Hot Start PCR Master Mix (Invitrogen) and 200 nM of each primer. Annealing temperatures were tested for each primer pair in gradient PCR to determine the one optimal for amplification.

Supplementary Table 7a shows the primers used for each gene target, the amplicon size, PCR reaction conditions, and NCBI accession numbers for sequences used primer design. PCRs were then performed with limited cycles of amplification to have a semi-quantitative comparison of transcript abundance between infection timepoints (by not reaching the maximum product amplification plateau). Amplification products were viewed using 1.5% agarose gel electrophoresis.

C315R transcript levels were assessed by qPCR, using housekeeping gene glyceraldehyde-3-phosphate dehydrogenase (GAPDH) expression was used for normalisation. Primer details and the qPCR amplification program are shown in

Supplementary Table 7b (GAPDH primers used for *Vero* cells were previously published by Melchjorsen et al., 2009 (105)). Primers were used at 250 nM concentration with Brilliant III Ultra-Fast SYBR® Green QPCR Master Mix (Agilent 600882), 1 µl cDNA in 20 µl (1:20) total reaction volumes, and qPCRs carried out in Mx3005P system (Agilent Technologies). Similar amplification efficiencies (97-102%) for all primers had been observed upon amplification of serially diluted cDNA samples, and the relative expression at each timepoint of infection was calculated using the formula 2^ΔCt^ (2^Ct_GAPDH-Ct_C315R^).

### Preparation of supernatant and cell lysis extracts for ELISA and Western blot detection of host proteins

Lung macrophage cultures from two donor outbred pigs (same cells used for CAGEseq) were prepared in 6-well plates. Approximately 1.5x 10^6^ cells were seeded per well with 3 ml medium (RPMI with penicillin/streptomycin and 10% pig serum) and incubated at 37 degrees 5% CO2 overnight. Cultures were washed once with culture medium to remove non-adherent cells and inoculated with MOI 5 of ASFV-Georgia 2007/1 (or left uninfected as control) and centrifuged 1h at 600xg 26 degrees (adsorption period). Supernatants from cell cultures were collected immediately after adsorption for obtaining the 0 hpi timepoint and stored at -70 degrees until analysis. Adherent cells were washed twice with cold DPBS (Sigma) and then lysed with 0.12 ml/well cold RIPA buffer (Thermo Scientific) supplemented with protease inhibitors (Halt Protease Inhibitor Cocktail, Thermo Scientific). For 5h and 16h timepoints, the inoculum was removed after adsorption, cells were washed twice in culture medium and returned to the incubator with fresh 3 ml medium per well for the specified times of infection. Supernatants and lysis volumes were collected similarly to the control. Supernatants were analysed for the presence of CCL2 (Porcine CCL2/MCP-1 ELISA Kit, ES2RB Invitrogen), CXCL8 (Quantikine® ELISA, Porcine IL-8/CXCL8 Immunoassay, P8000 R&D) and TNF-α (Quantikine® ELISA, Porcine TNF-α Immunoassay, PTA00 R&D) as recommended by the manufacturers. A volume of 25 µl each lysate was analysed in Western Blot for expression of ISG15 (anti-ISG15 antibody ab233071, Abcam; used at 1:1000 dilution), γ-Tubulin (anti-gamma Tubulin antibody ab11321, Abcam; used at 1:1000 dilution); and viral ASFV protein P30 (in-house mouse monoclonal antibody used at 1:500 dilution). Secondary antibodies used were Goat Anti-Rabbit IgG H&L (HRP) (ab205718, Abcam) and Goat Anti-Mouse Immunoglobulins/HRP (P0447, Dako) both at 1:2000 dilution. Western blot membranes were revealed using Pierce ECL Western Blotting Substrate (32106, Thermo Scientific). Band densities were quantified using ImageJ (Rasband, W.S., ImageJ, U. S. National Institutes of Health, Bethesda, Maryland, USA, https://imagej.nih.gov/ij/, 1997-2018).

### ASFV Promoter Motif Analysis

DESeq2 results were used to categorise ASFV genes into two simple sub-classes: early; 87 genes downregulated from early to late infection and late; the 78 upregulated from early to late infection. These characterised gene pTSSs were then pooled with the nORF pTSSs, and sequences upstream and downstream of the pTSS were extracted from the GRG genome in FASTA format using BEDtools. Sequences 35 bp upstream of and including the pTSSs were analysed using MEME software (http://meme-suite.org) (106), searching for 5 motifs with a maximum width of 20 nt and 27 nt, respectively (other settings at default). The input for MEME motif searches included sequences upstream of 134 early pTSSs (87 genes and 47 nORFs) for early promoter searching, while 234 late pTSSs (78 genes and 156 nORFs) were used to search for late promoters. For analysis of conserved motifs upstream of the five clusters described in Figure 6a-b, sequences were extracted in the same manner as above, but grouped according to their cluster. MEME motif searches were carried out for sequences in each cluster, searching for 3 motifs, 5-36 bp in length, with zero or one occurrence per sequence (‘zoops’ mode).

### Identification of TSSs by rapid amplification of cDNA ends -5’RACE

For 5’RACE of GRG genes DP146L, pNG4 and CP204L we designed the gene specific primers (GSP) shown in Supplementary Table 7c, and used the kit: “5’ RACE System for Rapid Amplification of cDNA Ends” (Invitrogen), according to manufacturer instructions. Briefly, 150 ng RNA from either 5 hpi or 16 hpi macrophages (one of the replicate RNA samples used for CAGE-seq) was used for cDNA synthesis with GSP1 primers, followed by degradation of the mRNA template with RNase Mix, and column purification of the cDNA. A homopolymeric tail was added to the cDNA 3’ends with Terminal deoxynucleotidyl transferase, which allowed PCR amplification with an “Abridged Anchor Primer” (AAP) from the 5’RACE kit and a nested GSP2 primer. A second PCR was performed over an aliquot of the previous, with 5’RACE “Abridged Universal amplification Primer” (AUAP), and an additional nested primer GSP3, except for pNG4 where GSP2 was re-used due to the small predicted size of the amplicon. Platinum™ Green Hot Start PCR Master Mix (Invitrogen) was used for PCR and products were run in 2% agarose gel electrophoresis (see

Supplementary Table 7c for expected sizes). Efficient recovery of cDNA from the purification column requires a product of at least 200 bases and therefore, due to the small predicted size of pNG4 transcripts its GSP1 primer was extended at the 5’ end with an irrelevant non-annealing sequence of extra 50 nt in order to create a longer recoverable product.

### CAGE-seq Analysis for the *Sus scrofa* Genome

Analyses of TSS-mapping, gene expression and motif searching with CAGE-seq reads mapped to the *Sus scrofa* 11.1 genome were carried out by DNAFORM (Yokohama, Kanagawa, Japan). The 5’ ends of CAGE-seq reads were utilised as input for the Reclu pipeline (107) with a cutoff of 0.1 RPM, and irreproducible discovery rate of 0.1. 37,159 total CAGE-seq peaks could be identified, of which around half (16, 720) match unique CAGE peaks previously identified by Roberts et al. (64) (i.e. within 100 nt of any of them). TSSs for 9,384 protein-coding genes (out of 21,288) were annotated de novo from the CAGE-defined TSSs (Supplementary Table 4).

Protein-coding genes with annotated TSSs (9,384 out of 21,288) were then subjected to differential expression analysis. CAGE-seq reads were summed up over all TSSs assigned to a gene and compared between two time points using edgeR (108) at maximum false discovery rate of 0.05. The full list of host genes with annotated promoters together with their estimated expression levels is provided in Supplementary Table 5. Gene set enrichment analysis was performed with the DAVID 6.8 Bioinformatics Resources (109), using best BLASTP (110) human hits (from the UniProt (111) reference human proteome). The 9,331 genes with human homologs were used as a background, and functional annotations of the four major expression response groups (late/early up-/down-regulated genes) were clustered in DAVID 6.8 using medium classification stringency. MEME motif searches were conducted for promoters of four differentially regulated subsets of host genes, as defined in Figure 11a. Promoters sequences were extended 1000 bp upstream and 200 bp downstream of TSSs, searched with MEME (max. 10 motifs, max. 100 bp long, on a given strand only, zero or one site per sequence, E < 0.01), and then compared against known vertebrate DNA motifs with Tomtom (p-value < 0.01).

### Data Availability

Raw sequencing data are available on the Sequence Read Archive (SRA) database under BioProject: PRJNA739166. This also includes CAGE-seq data aligned to the ASFV-GRG (FR682468.1 *Sus scrofa* (GCF_000003025.6) genomes (see methods above) in BAM format. Available for review via the link below: https://dataview.ncbi.nlm.nih.gov/object/PRJNA739166?reviewer=390lg85cohvh81llto5gr1d22n

## Acknowledgements

Research in the RNAP laboratory at UCL is funded by a Wellcome Investigator Award in Science ‘Mechanisms and Regulation of RNAP transcription’ to FW (WT 207446/Z/17/Z). GC is funded by the Wellcome Trust ISMB 4-year PhD programme ‘Macromolecular machines: interdisciplinary training grounds for structural, computational and chemical biology‘ (WT 108877/B/15/Z). Research in the ASFV group at The Pirbright Institute is funded by the Biotechnology and Biological Sciences Research Council (BBS/E/I/0007030 and BBS/E/I/0007031). The authors are grateful to all members of the RNAP lab and Tine Arnvig for critical reading of the manuscript.

## Notes

### Competing Interest Statement

The authors have declared no competing interest.

### Summary of Updates

Addition of further experiments, and text/figure edits to improve clarity.

